# Mirror-image mRNA display uncovers isoform-selective D-peptide macrocycles targeting a cryptic KRAS pocket

**DOI:** 10.64898/2026.05.20.726527

**Authors:** Matthew J. Mitcheltree, Nicole Boo, Nicolas Boyer, Zachary Z. Brown, Xiaomei Chai, Ruchia Duggal, Michael Garrigou, Robert P. Hayes, Jennifer M. Johnston, Hubert Josien, Brian M. Lacey, Shuhui Lim, Songnian Lin, Todd Mayhood, Haruhiko Ogawa, Peter Orth, Patrick C. Reid, Ryohei Shigeta, Aileen Soriano, Tatsuya Tomiyama, Gireedhar Venkatachalam, Yunpeng Zhou, David Jonathan Bennett, Anthony W. Partridge, Kaustav Biswas

## Abstract

Activating KRAS mutations drive millions of cancers diagnosed worldwide,^1^ yet for decades this oncoprotein was deemed “undruggable”, reflecting the challenge of discovering molecules capable of perturbing its complex biological functions, and of translating these discoveries into effective cancer therapeutics.^2^ Recent advances propelled by innovative screening have identified diverse modalities that bind at or near the switch-II pocket (SII-P) of RAS proteins, including molecular glues,^3^ macrocyclic peptides,^4^ fragment-derived small molecules,^5^ and approved therapies that covalently target KRAS^G12C^.^6,7^ Unfortunately, resistance to approved therapies has emerged,^8,9^ highlighting the need for molecules that engage new or underexploited binding sites on RAS oncoproteins with mechanisms complementary to established SII-P inhibitors.^10,11^ Here we show that mirror-image mRNA display^12^ enabled the discovery of all-D macrocyclic peptide ligands targeting a cryptic RAS back pocket (CRB-P).^13^ These ligands engage KRAS(OFF) and KRAS(ON) with equal affinity, exploit a single-residue difference among isoforms to bind KRAS selectively, and successfully inhibit oncogenic signaling in KRAS-mutant cells through a mechanism distinct from SII-P binders. Mirror-image screening directly afforded nanomolar peptide ligands stable toward cellular proteolysis and delivered probes targeting distinct epitopes not accessible by homochiral peptide-display methods. Together, these findings establish the CRB-P as a specifically druggable and mechanistically differentiated site on KRAS with potential for combination with emerging RAS-targeting therapies and substantiate mirror-image mRNA display as a strategy for discovering stable all-D macrocyclic peptides targeting previously inaccessible epitopes on challenging targets.

## Introduction

Activating mutations of the KRAS oncoprotein drive nearly a fifth of human cancers,^1^ underpinning a decades-long search for molecules capable of binding and inhibiting mutant KRAS with selectivity over its related isoforms, HRAS and NRAS.^2^ Innovative screening approaches have been essential to the discovery of such molecules, as the spheroid structure of the protein had for decades carried a reputation of being “undruggable”. Most ligands identified to date target the switch-II pocket (SII-P),^3,5,6,7^ emerging from covalent and fragment-based discovery efforts, with additional modalities—including macrocyclic peptides—largely converging on the same or nearby sites.^3,4,17,18^ While these efforts have culminated in the approval of covalent KRAS^G12C^ inhibitors and the clinical advancement of KRAS(OFF) conventional and KRAS(ON) tri-complex inhibitors, emerging resistance mechanisms such as adaptive feedback reactivation of the MAPK pathway and secondary KRAS mutations highlight the need for agents that operate through complementary mechanisms and are amenable to combination with existing therapies. Binders to distal pockets are also known,^14,15,16^ but rare, and often lack clear translational therapeutic potential. Recently, yeast display screening produced nanobody binders to a cryptic RAS back pocket (CRB-P);^13^ however, these binders lack isoform selectivity, exhibiting comparable affinity toward KRAS, HRAS, and NRAS. Notably, all known phage- and mRNA-display screens generating KRAS isoform-selective macrocyclic peptides to date have yielded SII-P binders,^4,17,18^ reinforcing the reputation of this pocket as perhaps the only site to which selective and functional KRAS ligands might be identified. Here we challenge this paradigm by applying mirror-image mRNA display^12^ to discover macrocyclic D-peptides exhibiting selective affinity for KRAS through isoform-specific interactions with the newly discovered CRB-P. These ligands inhibit oncogenic KRAS signaling through a mechanism distinct from SII-P binders and bind non-competitively with SII-P ligands, engaging both KRAS(ON) and KRAS(OFF) and establishing the CRB-P as a druggable site with potential for single-agent and combination strategies aimed at achieving more durable and efficacious pathway suppression. These findings support structure-based design of therapies exploiting this pharmacology and highlight mirror-image screening as an under-utilized strategy to discover new structure–function relationships for challenging biological targets.

### Discovery of D-peptide macrocycles with selective affinity for KRAS

We applied mirror-image mRNA display to uncover macrocyclic D-peptides targeting mutant KRAS. Specifically, we chemically synthesized the mirror-image form of KRAS^G12D^ (D-KRAS^G12D^) and refolded the purified, linear peptide in the presence of L-5’-guanylyl imidodiphosphate (L-GMPPNP) as previously described for D-KRAS^G12V^.^19^ We then employed the Peptide Discovery Platform System (PDPS^®^)^20,21,22^ to discover L-peptide macrocycles that bind to D-KRAS^G12D^ through iterative selection rounds against D-KRAS^G12D^(L-GMPPNP) (Figure 1a). We thus identified the L-configured enantiomer of **DMP-1** (Figure 1b) among a class of hit sequences enriched after next-generation sequencing (NGS, Extended Data Figure 2). SPR analysis demonstrated that the D-configured peptide bound recombinant KRAS with 1:1 stoichiometry and comparable affinity across wild-type and mutant forms in both GDP- (OFF) and D-GMPPCP-loaded (ON) states (Figure 1c). Comparable results were obtained when measuring the binding of **DMP-1** to KRAS^G12D^(GDP) and KRAS^G12V^(GDP) by isothermal titration calorimetry (ITC, Extended Data Figure 2c). Remarkably, and in contrast to previously described peptide and nanobody binders, **DMP-1** demonstrated pronounced isoform selectivity, featuring >60-fold greater affinity toward KRAS versus HRAS and NRAS (Figure 1b-c). We also observed selective affinity (∼30-fold) toward the 4B, versus 4A splice variant of KRAS^G12V^ (Figure 1b), prefiguring our discovery that the C-terminal helix α5 encoded by the alternative fourth exon of *KRAS* comprises part of the **DMP-1** binding site.

**Figure 1.**
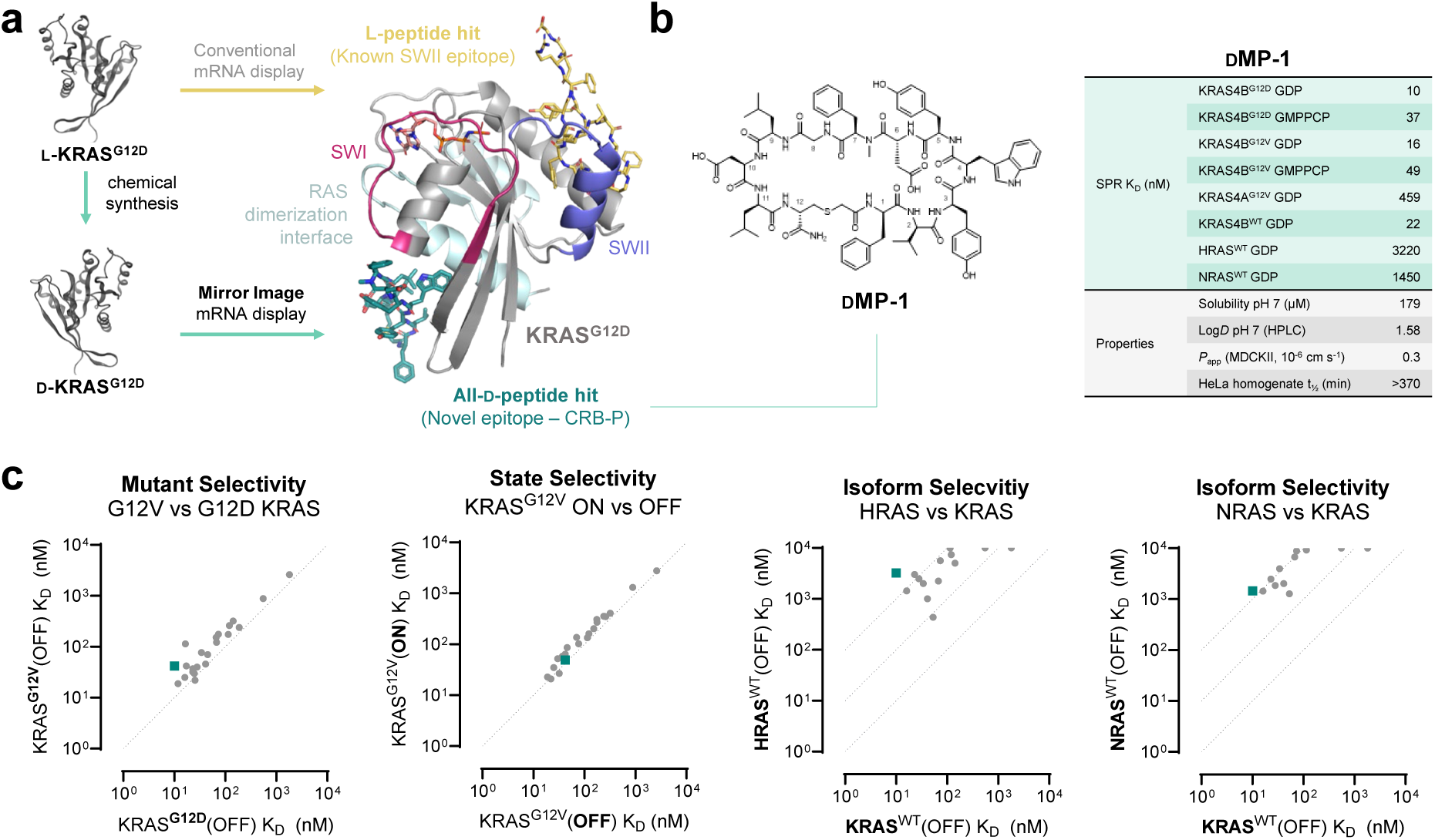
D-Peptide macrocycles selectively target common KRAS mutants. (a) Homochiral peptide screening historically has produced ligands to the well-studied SII-P (e.g., KD2 in gold, PDB: 6WGN). Our mirror-image screening approach instead uncovered ligands such as **DMP-1** which bind a cryptic back pocket of KRAS adjacent to the RAS dimerization interface. (b) **DMP-1** exhibits strong and selective affinity for KRAS irrespective of nucleotide loading state or common activating mutations. (c) **DMP-1** (teal square) and related congeners (gray dots) bind with equal affinities to KRAS^G12D^ and KRAS^G12V^, as well as to mutant KRAS in both its GMPPCP-loaded (ON) and GDP-loaded (OFF) states, while demonstrating 10- to 100-fold selectivity for KRAS over HRAS and NRAS isoforms.

These results prompted us to interrogate the biochemical activity of **DMP-1**. In a TR-FRET assay gauging displacement of a SII-P peptide binder,^23^ **DMP-1** elicited no effect (results not shown), demonstrating that **DMP-1** binds non-competitively with SII-P ligands. Nonetheless, **DMP-1** potently reduced the rate of SOS-mediated guanine nucleotide exchange (GNE) of various KRAS mutants (half-maximal effect concentrations ∼ 10-50 nM), whereas its activity against HRAS and NRAS was >100-fold lower, corresponding to differences in affinity measured biophysically among these proteins (Figures 2a-b). Notably, the maximal rate reductions observed for **DMP-1** and its analogues in these GNE experiments were approximately half of the theoretical maximum (SII-P ligands commonly achieve near-complete GNE inhibition), and so resembled the effects of a CRB-P nanobody ligand on spontaneous nucleotide exchange reported previously.^13^ In another departure from SII-P ligand pharmacology, **DMP-1** did not inhibit FRET between CRAF Ras-binding domain (RBD) and KRAS^G12D^, indicating that the peptide does not impair the interaction of these proteins (Figure 2c). Collectively, these results show that **DMP-1** and related analogs selectively engage KRAS through a binding site and mechanism distinct from SII-P ligands, modulating the rate of GEF-mediated nucleotide exchange without disrupting RAF association.

**Figure 2.**
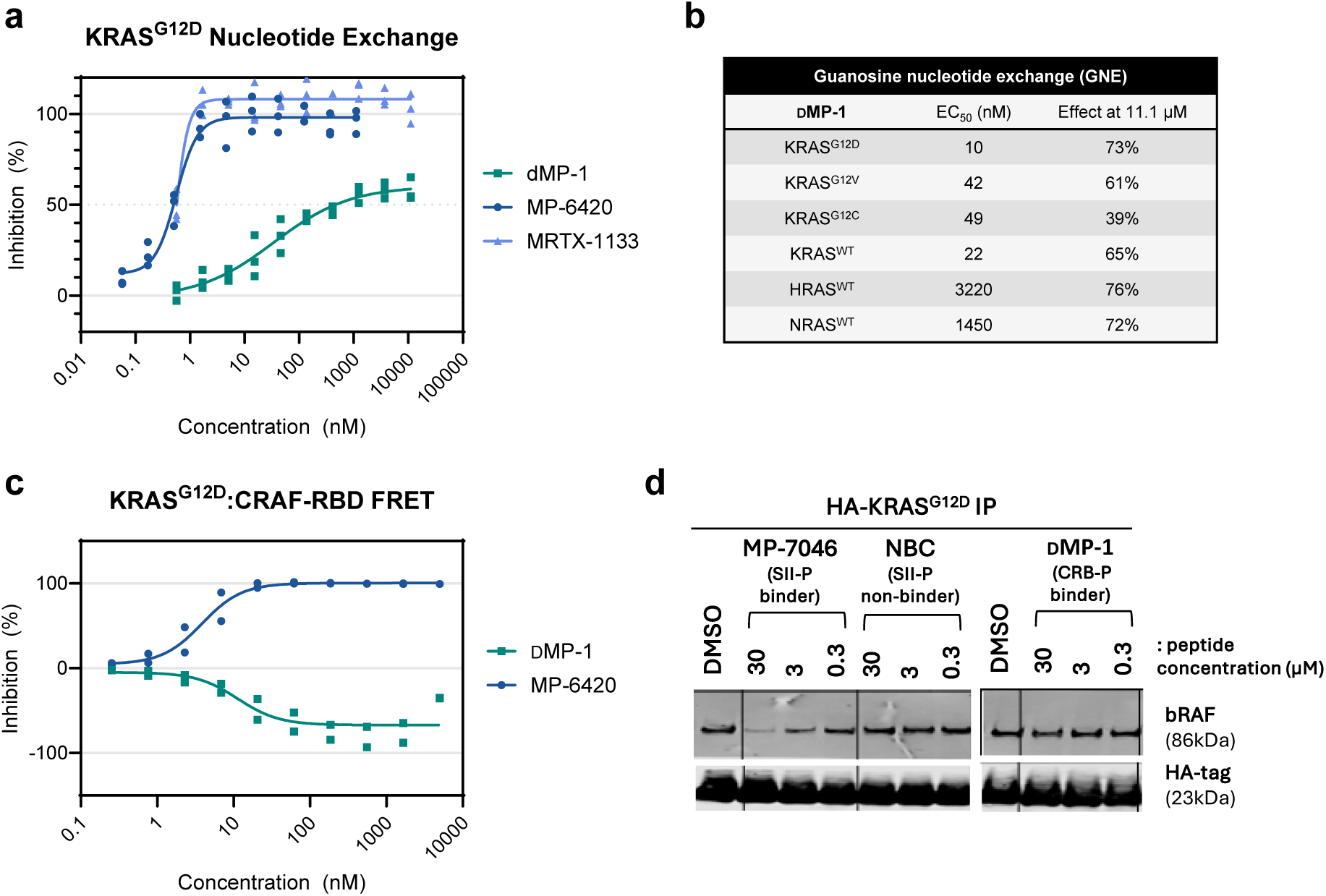
DMP-1 and SII-P ligands elicit distinct effects on KRAS GNE and RAF association. (a) Whereas SII-P ligands MP-6420 and MRTX-1133 achieve complete inhibition of KRAS^G12D^ SOS-mediated GNE, **DMP-1** reduces this rate by roughly half at maximal effect. (b) Half-maximal inhibitory concentrations and maximal response depth of **DMP-1** in SOS-mediated GNE of various KRAS mutants and RAS isoforms. (c) SII-P ligand MP-6420 and CRB-P ligand **DMP-1** exhibit divergent effects in KRAS^G12D^:CRAF-RBD FRET, as **DMP-1** dose-dependently enhances FRET signal between the two protein partners, plotted here as negative inhibition. (d) Co-immunoprecipitation of HA-tagged KRAS^G12D^ demonstrated dose dependent disruption of BRAF with SII-P ligand MP-7046, but not with **DMP-1** or **DMP-NBC** (non-binding control).

### Structure illuminates novel binding site and basis for selectivity

To understand the unique and isoform-specific pharmacology of **DMP-1**, we mapped the peptide binding epitope of KRAS^G12D^(GDP) by NMR and subsequently solved a crystal structure of the peptide bound to KRAS^G12D^(GMPPNP) (Figure 3 and Extended Data Figure 3). We found **DMP-1** binds the recently reported CRB-P, a site for which the identification of selective binders has been deemed improbable, owing to high residue homology among RAS isoforms.^13^ Upon binding the CRB-P, we observed that **DMP-1** adopts a β-hairpin structure, with four intramolecular hydrogen bonds and a trans-annular side-chain interaction (H-bonding between DTyr5 and DAsp10) supporting the bound conformation. The hairpin structure of the backbone of **DMP-1** neatly interleaves with KRAS β2 through a network of backbone hydrogen bonding with Val45 and Asp47, and a further backbone–sidechain interaction with Asp47. A charged interaction between the sidechains of ligand residue DAsp6 and KRAS residue Lys42 completes this set of interactions to effectively extend the β2-β3 sheet, while cryptic back-pocket residue Tyr157 re-orients to accommodate **DMP-1** residue DTrp4. This pivotal ligand residue—DTrp4, tellingly enriched among hits in mRNA display sequencing data (Extended Data Figure 2a)—simultaneously interacts with the lipophilic, cryptic pocket revealed by re-orientation of Tyr157 to anchor the peptide against the surface, while forming a direct H-bonding interaction with the α5 residue Asp153 of KRAS splice variant 4B. It is to this interaction with Asp153 that we attribute **DMP-1**’s remarkable isoform selectivity, as this residue is unique to KRAS4B; HRAS, NRAS, and KRAS4A bear larger and more flexible glutamate residues at the corresponding site and thus present less optimal interactions with **DMP-1**. Indeed, free-energy perturbation simulations for the alchemical transformation of KRAS4B Asp153→Glu predict a free-energy difference ΔΔG_T=300K_ = 2.4 ± 0.5 kcal mol^−1^, demonstrating this residue alone can account for the ∼30- to 100-fold selective affinity **DMP-1** features toward KRAS4B relative to HRAS, NRAS, and KRAS4A (Figure 1b).

**Figure 3.**
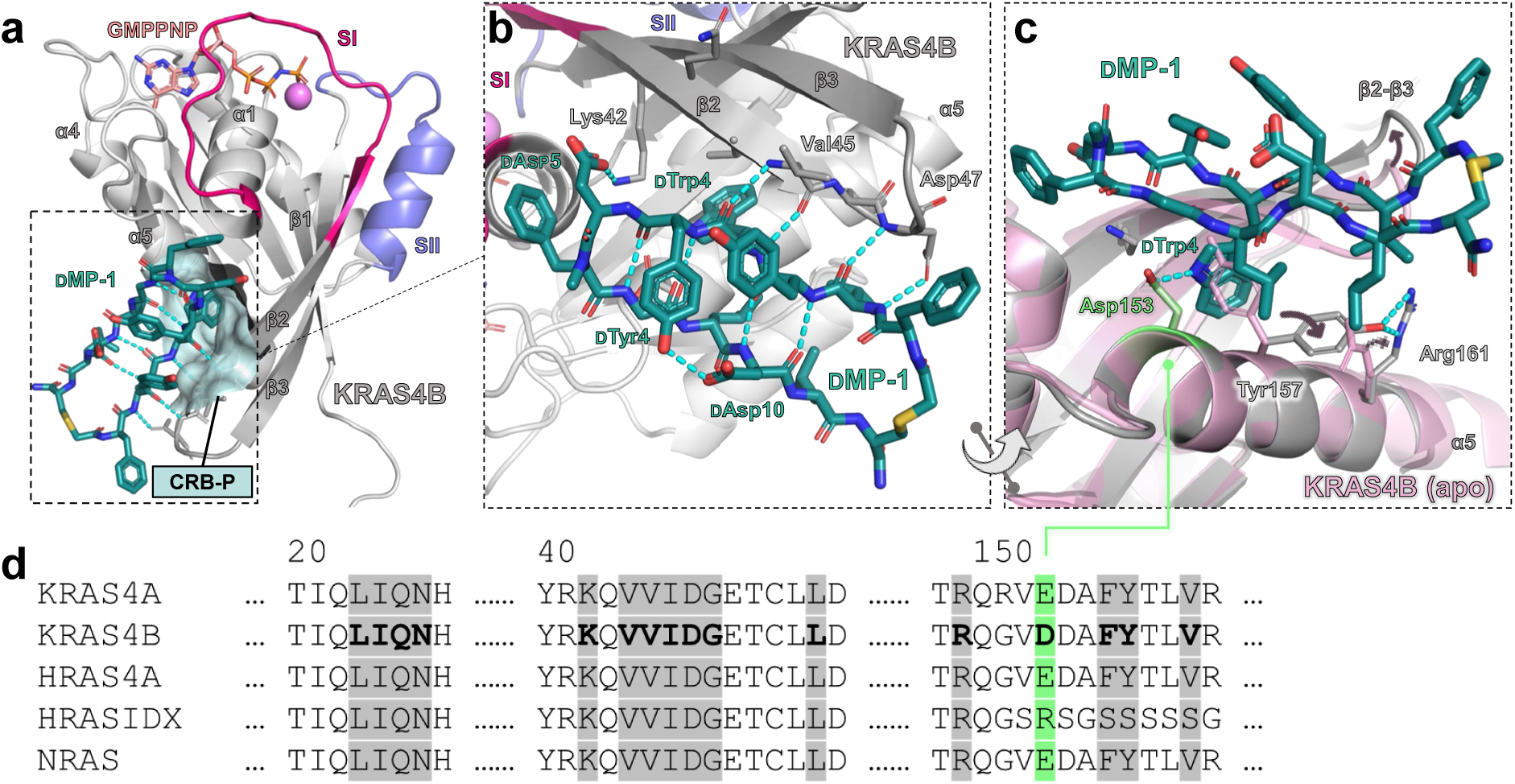
Structural basis for DMP-1’s selective affinity toward KRAS4B. (a) X-ray crystallographic structure of **DMP-1** (teal sticks) bound to KRAS4B^G12D^(GMPPNP) (grey protein, salmon nucleotide) reveals the peptide binds a cryptic pocket (CRB-P, teal surface) distal from switch I (SI, crimson) and switch II (SII, cornflower) regions more commonly targeted by RAS ligands. (b) **DMP-1** binds the CRB-P in a β-hairpin conformation, extending the β3-β2 sheet of KRAS through backbone and side-chain H-bonding, and projecting lipophilic sidechains, notably including DTrp4, into a pocket abutting KRAS helix α5. (c) Structural overlay with unliganded KRAS4B (6EPL, rose)^32^ illustrates induced fit involving side-chain displacement of α5 residues Tyr157 and Arg161 to accommodate **DMP-1**. The indole ring of ligand residue DTrp4 forms an anchoring H-bond with KRAS4B residue Asp153 (lime green). (d) Aligned sequences of RAS isoforms illustrate the single-residue structural basis for **DMP-1**’s observed isoform selectivity. KRAS4B residues observed crystallographically to contact **DMP-1** are bolded, and corresponding positions are highlighted across sequences. Asp153 (lime green) is the sole residue contacting **DMP-1** to differ among these sequences.

### DMP-1 engages mutant KRAS in cells to inhibit downstream signaling

Having established that **DMP-1** binds selectively in the CRB-P of KRAS mutants, we sought to understand the potential of CRB-P binders to perturb mutant KRAS signaling in cells by modulating downstream signaling via the RAS-MAPK-ERK pathway. In HeLa cell homogenate, **DMP-1** exhibited excellent proteolytic stability (t_½_ > 6 h), consistent with expectations for a macrocycle comprising exclusively D-configured amino acids and supporting its use as a cellular tool compound. Hence, we first evaluated whether **DMP-1** could engage KRAS in cells. A biotinylated affinity probe (**DMP-BTN**, Extended Data Figure 4) successfully pulled down membrane-bound KRAS^G12D^ from AsPC-1 membrane patch isolates, and this interaction was dose-dependently competed by non-biotinylated **DMP-1**, but not by a non-binding control (**DMP-NBC**) in which residues DVal2 and DTrp4, crystallographically revealed to form vital interactions with KRAS, were transposed (Figure 4a). In cellular thermal shift assays (CETSA),^24^ **DMP-1** conferred thermal stabilization of KRAS^G12C^ in intact cells. This effect was enhanced by co-incubation with digitonin, introduced to boost intracellular exposure of the otherwise poorly permeable peptide (Figure 4b). Indeed, consistent with its high molecular weight and exposed polarity, **DMP-1** exhibited poor passive permeability (MDCKII P_app_ = 0.3 × 10^−6^ cm s^−1^) and no effect on ERK phosphorylation in intact AsPC-1 cells (Figure 4c). To overcome this limitation, we pursued two complementary delivery strategies. First, we designed a cell-penetrating variant (**DMP-CPP**) incorporating six contiguous DArg residues at the N-terminus of **DMP-1**. This peptide achieved complete inhibition of ERK phosphorylation in AsPC-1 cells with a half-maximal effect concentration (EC_50_ = 2.8 μM) typical of KRAS-inhibitory peptides relying on active transport for cellular uptake (Figure 4c).^17,23,25^ Second, digitonin-mediated membrane permeabilization enabled **DMP-1** itself to inhibit ERK phosphorylation in AsPC-1 (KRAS^G12D^), Panc 08.13 (KRAS^G12D^), and SK-CO1 (KRAS^G12V^) cells with micromolar potency, while nonbinding control **DMP-NBC** had no effect (Figure 4d-e). Importantly, **DMP-1** showed no activity in digitonin pre-treated A375 cells harboring the BRAF^V600E^ mutation, which activates the MAPK pathway independently of KRAS, confirming that pERK inhibition by CRB-P-targeting peptides is mediated through direct KRAS engagement (Figure 4e). Finally, to probe the mechanism by which CRB-P ligands inhibit MAPK signaling, we examined the effect of **DMP-1** on the KRAS-RAF interaction in cells. In contrast to SII-P binders such as **MP-7046**, which reduced co-immunoprecipitation of BRAF with HA-tagged KRAS^G12D^ from cell lysates, **DMP-1** had no effect on KRAS-BRAF association under identical conditions (Figure 2d), mirroring our biochemical observation that **DMP-1** does not inhibit KRAS–CRAF RBD FRET (Figure 2c). Taken together, these findings demonstrate that CRB-P ligands inhibit oncogenic KRAS-MAPK signaling across multiple KRAS-mutant cellular contexts through a mechanism that is dependent on KRAS engagement but distinct from direct disruption of the KRAS-RAF interaction.

**Figure 4.**
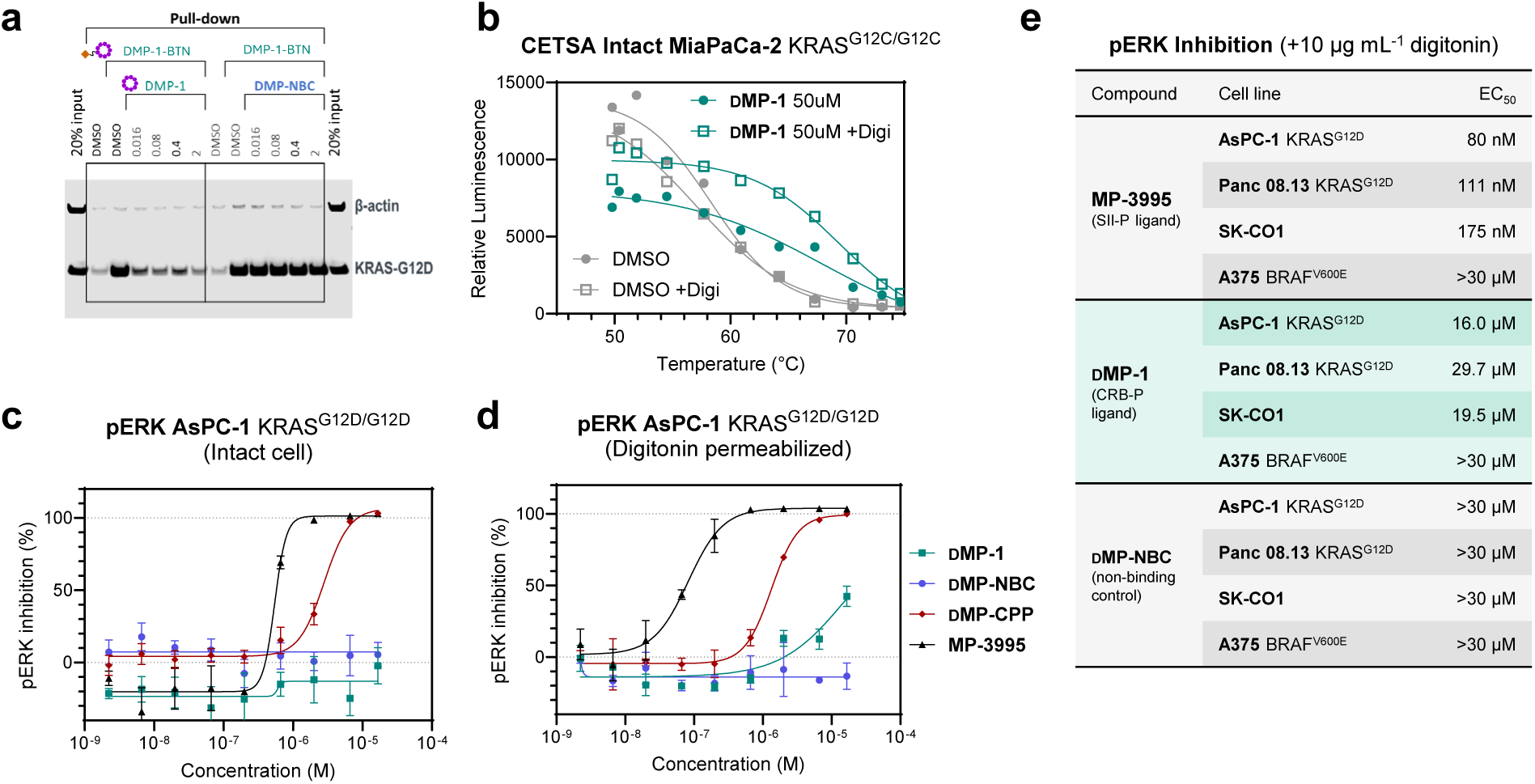
DMP-1 engages mutant KRAS in cells to inhibit downstream signaling. (a) Streptavidin pull-down of AsPC-1 cell lysate followed by KRAS immunoblotting shows robust enrichment of KRAS**^G12D^** by biotin-tagged probe **DMP-BTN**, competitively displaced by coincubation with untagged DMP-1, but not by nonbinding control peptide **DMP-NBC**. (b) **DMP-1** thermally stabilizes KRAS^G12C^ in intact MiaPaCa-2 cells treated with 10 μg mL^−1^ digitonin. Thermal stability was observed compared to DMSO control with ∼9 °C increase in melting temperature (T_m_). (c) Cell-penetrating peptides derived from SII-P (**MP-3995**) and CRB-P ligands (**DMP-CPP**) inhibit ERK phosphorylation in AsPC-1 cells, while impermeable peptides **DMP-1** and **DMP-NBC** do not. (d) Cell permeabilization with digitonin (10 μg mL^−1^) enhances the functional activity of **DMP-1**, but not of cell-penetrating peptide analog **DMP-CPP** or of non-binder control **DMP-NBC**. Data in **c** and **d** are mean ± s.e.m. of triplicate wells. (e) Digitonin-permeabilized cells harboring activating KRAS mutations, but not downstream mutations, exhibit dose-dependent inhibition of ERK phosphorylation upon treatment with SII-P and CRB-P ligands (**MP-3995** and **DMP-1**, respectively), but not upon treatment with **DMP-NBC**.

## Discussion

Without epitope-blocking or other means of biasing the discovery of hits targeting new epitopes, mirror-image mRNA display uncovered ligands to a cryptic binding site on KRAS that has eluded previous screens of small-molecule and naturally configured peptide libraries. This finding highlights potential for mirror-image display to discover D-peptide macrocycles targeting previously unknown or inaccessible sites on other challenging targets, with attendant possibilities for new mechanisms of action and isoform selectivity as we describe here.

mRNA display libraries, while very large in sequence space (10^12^-10^15^), are constrained in conformation space by the uniform constitution and configuration of their L-α-aminoacyl backbone, such that their diversity can be conceived as emerging through a high degree of side-chain variation laid upon a narrower range of three-dimensional scaffolds. Thus, we speculate that screening the same peptide-mRNA library against two enantiomeric forms of the same target—in effect doubling the chemical space described by peptide backbone conformations—may have a greater impact on hit identification than a doubling of the size of the homochiral peptide library itself. Illustrating this point, the central pharmacophoric interactions underpinning **DMP-1**’s affinity and selectivity towards KRAS appear to be uniquely accessible to D-, and not L-peptide macrocycles: Only a non-naturally configured peptide can extend the β2-β3 sheet of KRAS through backbone H-bonding interactions, position the selectivity-determining residue, DTrp, within the cryptic pocket, and avoid side-chain clashes with adjacent KRAS residues (e.g., Val44, Ile46). Whereas synthetic access to D-proteins historically has presented significant challenges, we note that recent methodological advances in solid-phase protein synthesis may expand the scope of proteins suited to mirror-image display screens.^26^

Our findings indicate that CRB-P ligands such as **DMP-1** inhibit KRAS function through a mechanism fundamentally distinct from SII-P binders, and parallel observations from studies of dimerization-impaired KRAS mutants. Specifically, Ambrogio and colleagues have shown that D154Q mutation impairs KRAS dimerization without impacting GEF or GAP sensitivity or RAF binding, and employed this mutant to show that dimerization of oncogenic KRAS is essential to downstream signaling.^27^ We find that **DMP-1** binds the same interface, engaging D153 and adjacent residues, and similarly does not impair RAF binding. This interface is largely identical to the previously reported KRAS α4-α5 dimerization surface, suggesting that CRB-P ligands achieve pathway inhibition by disrupting KRAS dimerization—a process required for downstream signaling, but not for RAF recruitment (Extended Data Figure 5). One notable distinction is that **DMP-1** partially inhibits SOS-mediated nucleotide exchange, whereas D154Q does not. While previously reported RAS-SOS co-crystal structures show D154 does not interact with SOS directly,^28^ our structure shows that **DMP-1** occludes KRAS binding to the activating allosteric site (but not the catalytic site) of SOS (Extended Data Figure 5b), offering a plausible explanation for this difference. Together, our data support a model in which CRB-P ligands act to suppress oncogenic KRAS signaling primarily through disruption of KRAS dimerization, and define a mechanistically distinct mode of KRAS inhibition. Because CRB-P ligands bind non-competitively with SII-P ligands, this mechanism of action is inherently compatible with existing and emerging KRAS-targeting therapies, raising the prospect of combination strategies that simultaneously target multiple functional sites on KRAS to achieve deeper or more durable pathway suppression with less propensity toward resistance.

The selectivity of **DMP-1** and its congeners for KRAS4B over KRAS4A, HRAS, and NRAS carries significant translational implications. KRAS4B is the predominant splice variant expressed in most human tissues and is most strongly implicated in oncogenic signaling across KRAS-driven malignancies, including pancreatic ductal adenocarcinoma, colorectal cancer, and non-small cell lung cancer.^29^ Therapeutics derived from these peptides that retain KRAS4B selectivity may therefore achieve robust efficacy while minimizing effects on KRAS4A-dependent processes in normal tissues. On the other hand, recent work also highlights emerging evidence of 4A splice variant signaling in KRAS cancer biology, the significance of which has historically been under-appreciated,^30,31^ and which may be more precisely interrogated using 4B-selective probes derived from the ligands we describe here. Indeed, the structural basis we report for targeting of KRAS4B by **DMP-1** provides a clear framework for the rational discovery and optimization of selective CRB-P ligands with the potential both to unlock deeper understanding of KRAS cancer biology, and to deliver translational therapies amenable to combination with SII-P-targeting medicines in development today. As clinical experience with KRAS inhibitors continues to uncover the challenges of acquired resistance driven by adaptive feedback reactivation, secondary KRAS mutations, and activation of bypass signaling pathways, mechanistically orthogonal agents such as CRB-P ligands are well positioned to serve as key components of combination regimens designed to address these resistance mechanisms.^11^ Consistent with their size and polarity, these 12-mer macrocyclic peptides exhibit poor passive permeability, a common feature of this modality; however, their robust activity upon facilitated intracellular delivery suggests that advances in delivery strategies or property optimization may enable broader application of CRB-P-targeting ligands as both biological probes and therapeutic agents.

## Extended Data Display Items

**Extended Data Figure 1.**
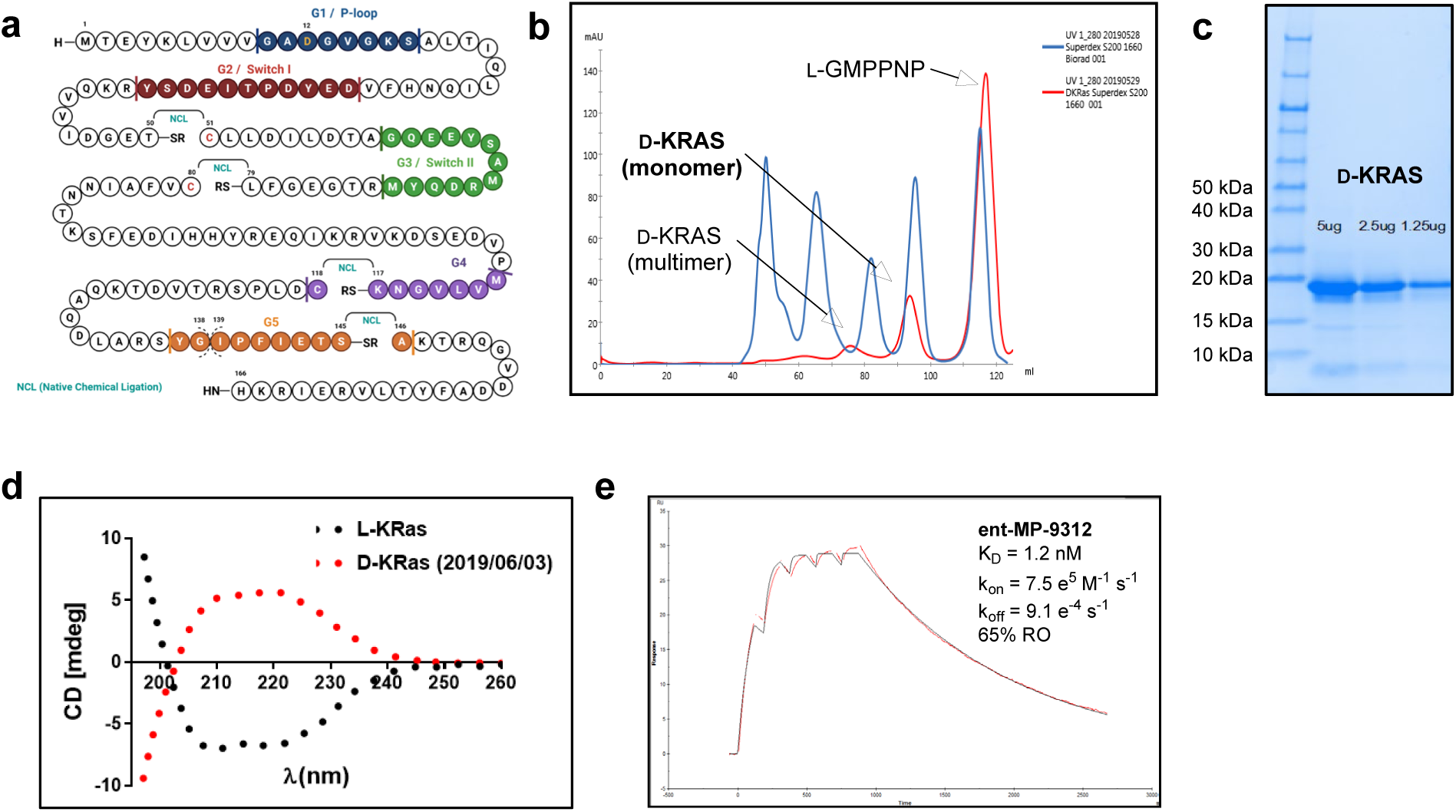
Chemical Synthesis of all-D KRAS^G12D^, purity assessment and validation of secondary and tertiary structure folding in the presence of L-GMPPNP. (a) Primary structure of KRAS^G12D^ (splice variant 4B, residues 1-166) and retrosynthetic strategy for native chemical ligation. Characterization of D-KRAS^G12D^ refolded in the presence of L-GMPPNP by analytical size-exclusion chromatography (b) and SDS-PAGE (c) confirmed the isolation of monomeric KRAS protein. (d) Circular dichroism (CD) analysis of folded and purified L-KRAS^G12D^ and D-KRAS^G12D^ with mirror-image protein exhibits an inverted CD spectrum compared to the naturally configured protein. (e) SPR sensorgram illustrating immobilized mirror-image protein binds ent-MP-9312, the D-configured enantiomer of known SII-P peptide ligand MP-9312.^23^

**Extended Data Figure 2.**
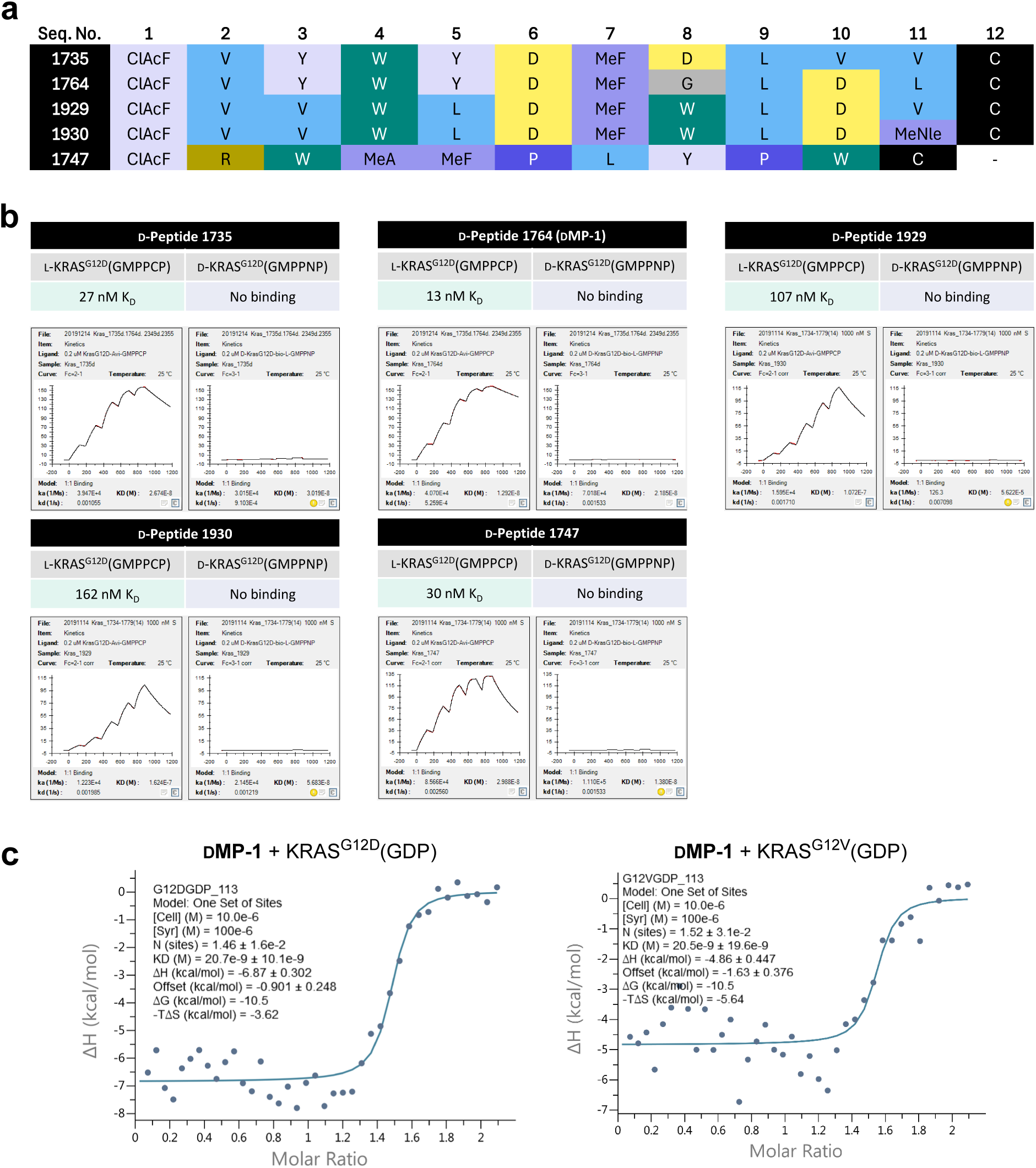
D-Peptide mRNA display hits selectively bind L-, not D-KRAS. (a) Representative sequences enriched after affinity selection via mirror-image mRNA display against D-KRAS^G12D^, including sequence number 1764 corresponding to **DMP-1**. (b) SPR sensorgrams of D-configured macrocyclic peptides and measured affinities for D- and L-configured KRAS^G12D^, confirming specificity of binding to the mirror-image protein form. (c) Isothermal titration calorimetry confirms affinity and 1:1 binding stoichiometry of **DMP-1** toward KRAS^G12D^(GDP) and KRAS^G12V^(GDP).

**Extended Data Figure 3.**
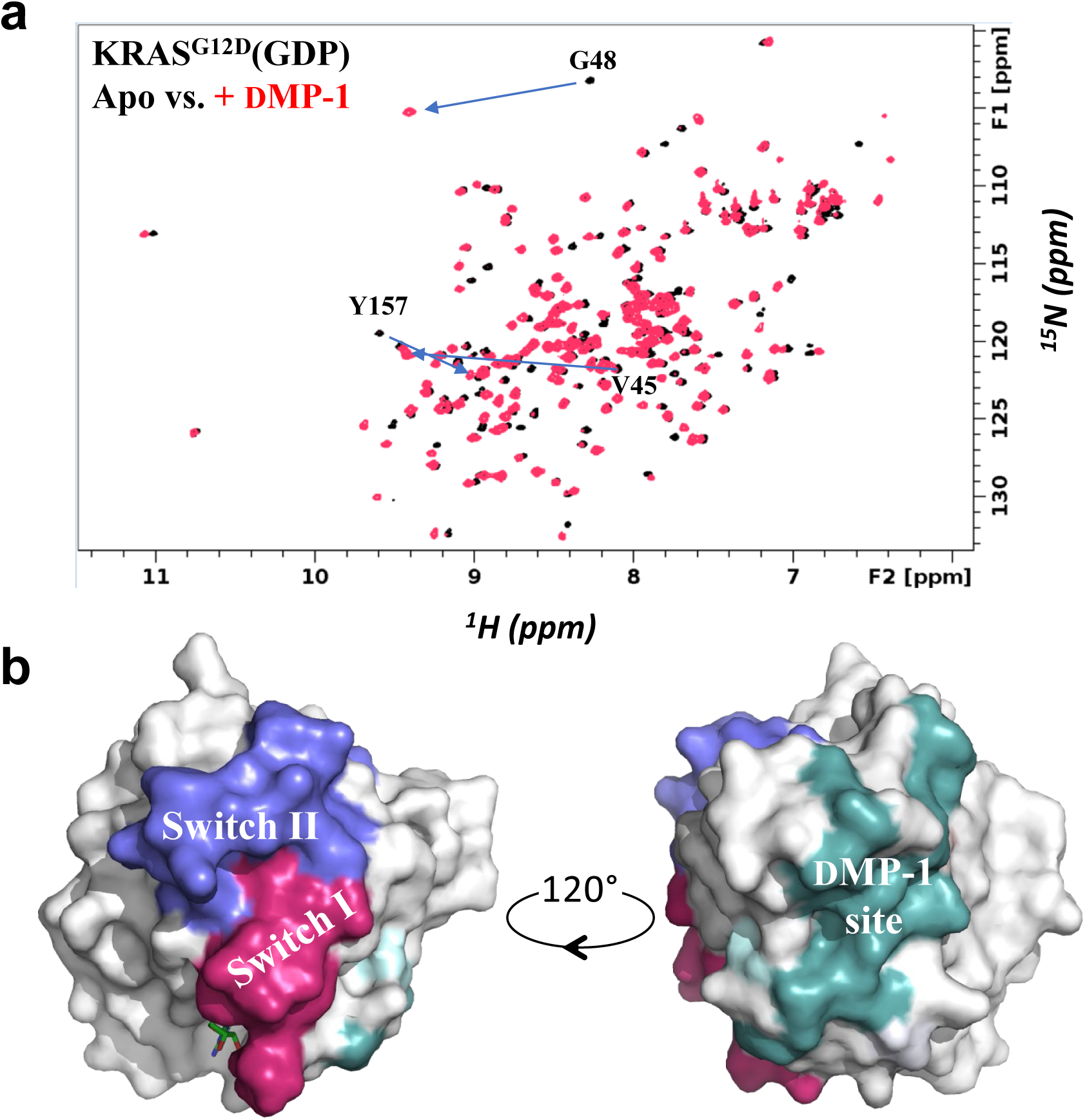
Mapping of the DMP-1 binding site by NMR. (a) ^1^H-^15^N HSQC spectrum of KRAS^G12D^(GDP) shows chemical shift perturbations induced by **DMP-1**. Particularly pronounced perturbations of Val45 (β2), Gly48 (β2-β3), and Tyr157 (α5) resonances are noted with blue arrows. (b) Mapping chemical shift perturbations on KRAS (PDB 4EPR) shows **DMP-1** binds a site on KRAS (teal) opposite to the Switch I and Switch II regions (crimson and blue, respectively)

**Extended Data Figure 4.**
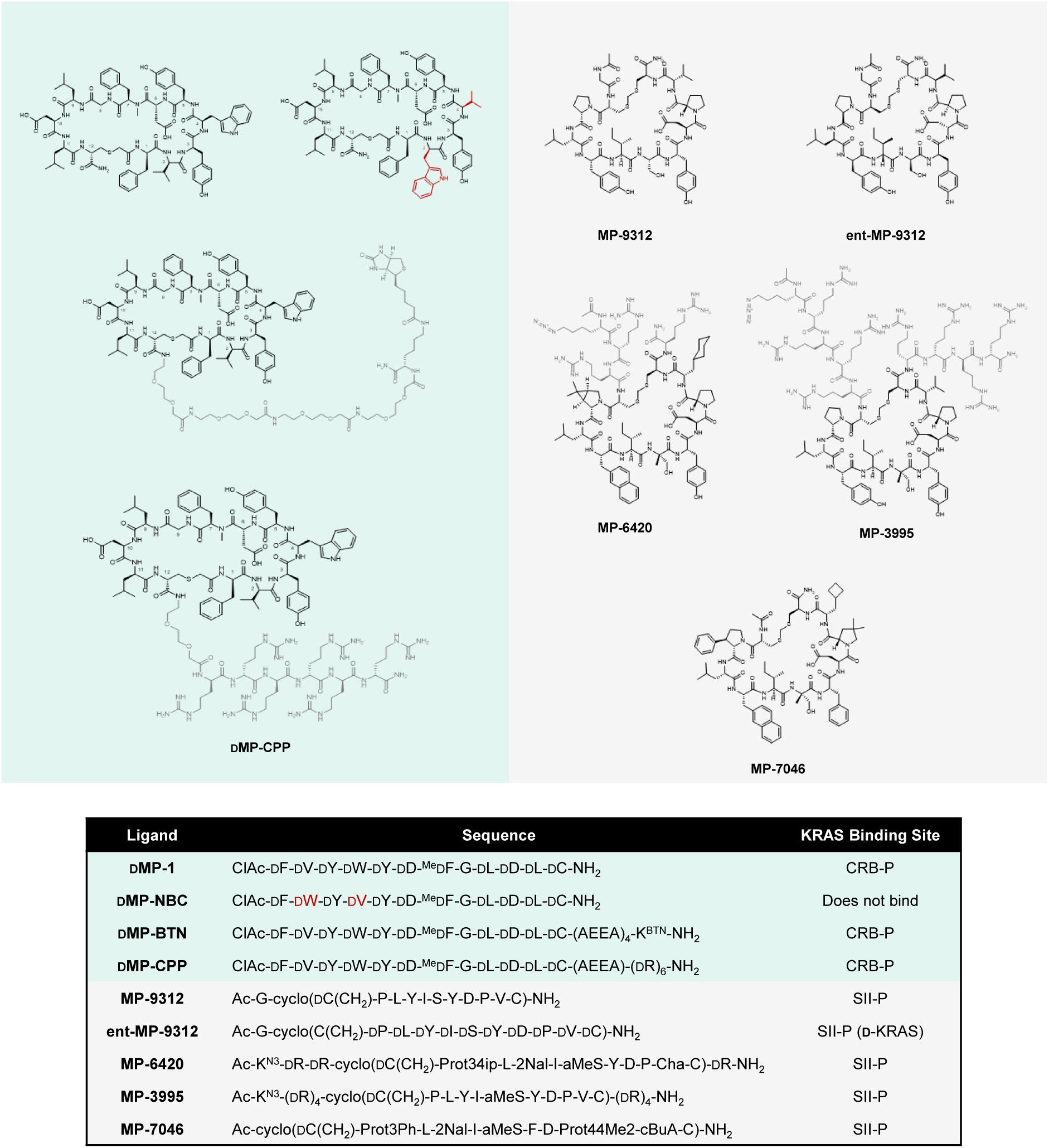
Structures, sequences, and binding sites of KRAS-targeting peptides.

**Extended Data Figure 5.**
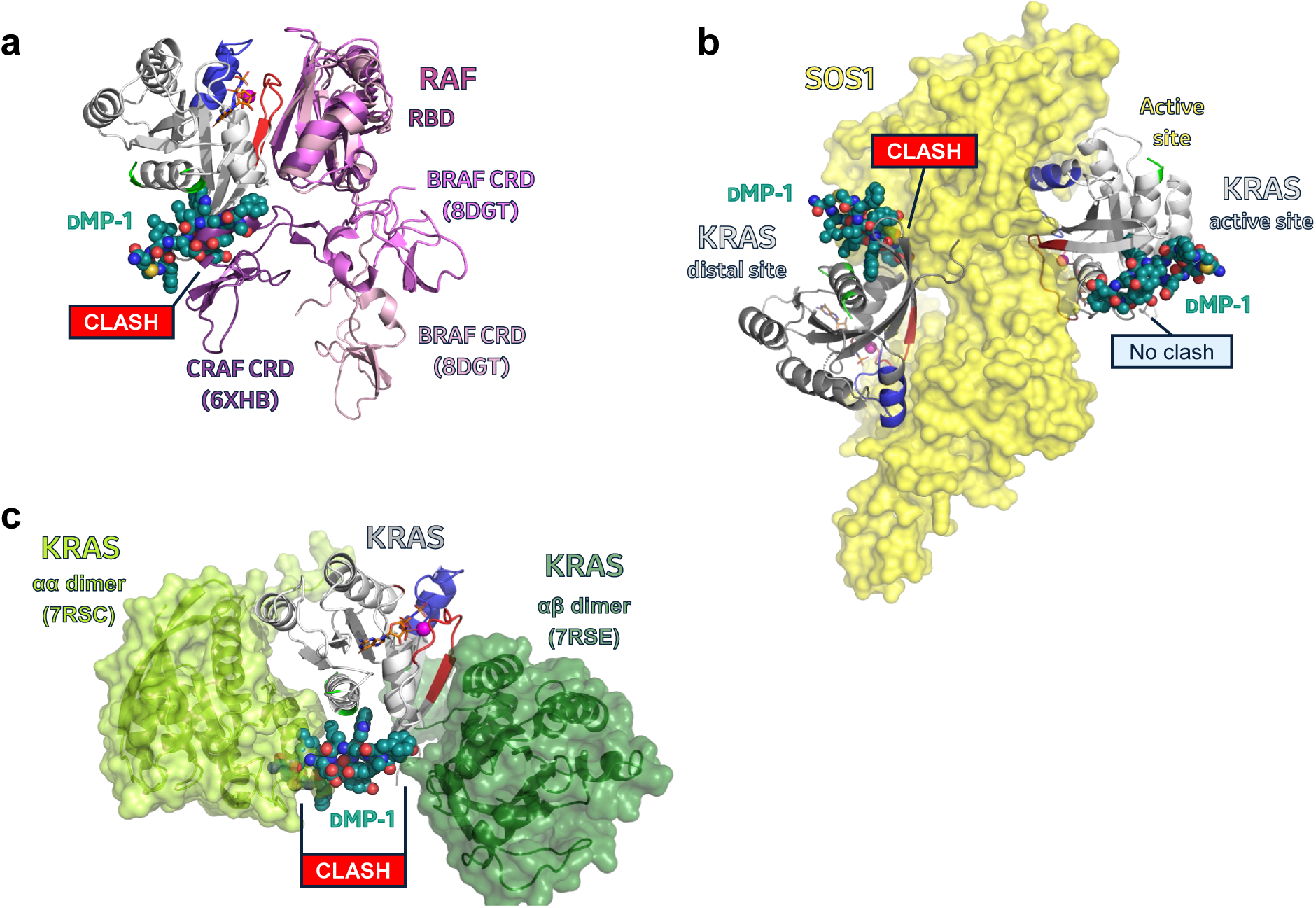
Overlay of DMP-1 binding site with KRAS interactor interfaces. (a) Superposition with reported KRAS–RAF structures illustrates that **DMP-1** binding does not occlude RAF RBD association. BRAF–KRAS ‘front’ (pink, PDB 8DGT) and ‘up’ (light pink, PDB 8DGS) structures extracted from MEK–BRAF–14-3-3–KRAS recruitment complex;^33^ CRAF–KRAS structure (purple, PDB 6XHB) extracted from NMR-nanodisc structure.^34^ (b) Overlay with reported RAS–SOS–RAS tri-protein complex shows **DMP-1**-bound KRAS appears competent toward active-site binding, but clashes at the distal site responsible for allosteric acceleration of SOS1-stimulated RAS activation via nucleotide exchange. (c) **DMP-1**-bound KRAS appears incompetent toward dimerization by the specific and symmetrical α–α interface (lime green, PDB 7RSC) illuminated by nanodisc NMR,^35^ while nonspecific α–β dimerization may also be affected (forest green, PDB 7RSE).

## AUTHOR CONTRIBUTIONS

M.J.M., A.W.P., N. Boo, and J.M.J. wrote the manuscript.

T.M. prepared proteins for screening and biochemical and biophysical analysis.

T.T. designed and supervised the mRNA display screen; R.S. performed the screen.

A.S. and H.O. performed SPR experiments.

N. Boo, R.D., B.M.L., S.L., G.V., and X.C. performed biochemical and cell biology experiments.

N. Boyer, Z.Z.B., M.G., H.J. designed and synthesized the compounds.

Y.Z. performed and interpreted NMR experiments.

P.O. and R.P.H. determined the X-ray structure of the peptide–KRAS complex.

J.M.J. and M.J.M. performed computational modeling.

H.O., P.C.R., R.S., and T.T. performed mRNA display screening and early hit validation.

A.W.P., K.B., S. Lin, P.C.R. and D.J.B. supervised the research.

All authors interpreted the results, reviewed and approved the manuscript.

## ACKNOWLEDGEMENTS

The authors gratefully acknowledge Yen Ting Chen, Andrew Le, Lisa O’Callaghan, Michael Eddins, and My Sam Mansueto for their contributions to this manuscript and thank Anandan Palani for his help in procuring material used to prepare mirror-image protein.

This research used resources at the Industrial Macromolecular Crystallography Association Collaborative Access Team (IMCA-CAT) beamline 17-ID, supported by the companies of the Industrial Macromolecular Crystallography Association. This research was performed on APS beam time award (DOI: https://doi.org/10.46936/APS-192278/60015810) from the Advanced Photon Source, a U.S. Department of Energy (DOE) Office of Science user facility operated for the DOE Office of Science by Argonne National Laboratory under Contract No. DE-AC02-06CH11357.

## ACCESSION NUMBERS

Protein Data Bank: Atomic coordinates for the **DMP-1**–KRAS^G12D^(GMPPNP) structure have been deposited under accession code 12XR.

## METHODS

### Synthesis of biotinylated D-KRAS^G12D^ via native chemical ligation

D-KRAS4B^G12D^ protein C-terminally tagged with biotin was prepared via Fmoc solid-phase peptide synthesis and subsequent native chemical ligation following the procedures described by Levinson *et al*. in their synthesis of D-KRAS^G12V^.^19^ Protein was refolded in the presence of L-GMPPNP, purified via size-exclusion chromatography, analyzed by circular dichroism (CD) spectroscopy, and quantified by absorbance at 280 nm according to methods described in the original report.

### Affinity selection against D-KRAS^G12D^

From the initial library theoretically consisting of more than 50 billion unique sequences, binders toward D-KRAS4B^G12D^ were enriched using biotinylated D-KRAS4B^G12D^(L-GMPPNP) immobilized on streptavidin-coated magnetic beads (Dynabeads M280; Thermo Fisher Scientific catalog no. 11205D) via iterative selection rounds. The enriched sequences were analyzed by next-generation sequencing, and highly frequent sequences were selected for chemical synthesis to afford D-configured peptide macrocycle ligands to L-KRAS.

### Protein production

Human KRAS4B (residues 2-169), KRAS4A (residues 2-169), NRAS (residues 2-172) and HRAS (residues 2-166) variants were synthesized and subcloned into the pET28a vector (Novagen; Millipore Sigma) with a N-terminal 6 × His tag, TEV cleavage site, followed by a SUMO tag and Avi tag. An additional KRAS4B (2-169) construct was designed to remove the Avi tag leaving native KRAS4B amino acids for BioNMR studies.

Plasmids were transformed into *E.coli* BL21(DE3) cells (NEB) followed by single colony selection on Luria-Bertani (LB) agar plates (Teknova) containing 50 μg mL^−1^ kanamycin. LB media (Teknova) containing 50 μg mL^−1^ kanamycin were inoculated with a single colony and grown overnight at 30 °C. Overnight cultures were diluted and grown at 37 °C until OD_600_ reached 0.8. Induction was initiated by adding 0.8 mM IPTG for 4 hrs until the cell density at OD_600_ reached 3.0. All Avi tag construct expressions were performed at 37 °C in LB media. Expression of double labeled (U-^15^N, U-^13^C) KRAS4B (2-169) for BioNMR studies was carried out in M9 minimal media using ^15^N NH_4_Cl (1 g L^−1^) and ^13^C glucose (4 g L^−1^) as the nitrogen and carbon sources, respectively. BioExpress (U-^15^N, U-^13^C) at 0.1X concentration was added to media to boost expression levels. Isotopes were purchased from Cambridge Isotope Laboratories.

Induced cell pellets were resuspended and lysed in Buffer A; 40 mM HEPES pH 8.0, 300 mM NaCl, 20 mM imidazole, 2 mM beta-mercaptoethanol using dounce homogenization followed by sonication (VibraCell VCX 750; Sonics). Cell debris was removed by centrifugation for 1 hr at 40,000 *g*. Clarified lysates were applied to a His-Trap Crude Fast Flow column (Cytiva) and eluted with Buffer A including 500 mM imidazole. Peak fractions were pooled and desalted into Buffer A for SUMO protease cleavage reaction (1 mg SUMO protease/40 mg RAS) carried out overnight at 4°C. RAS proteins were further purified using His-Trap Fast Flow to remove 6 × His-SUMO tag, SUMO protease and other contaminates in previously described buffers. Avi tag constructs were biotinylated using BirA ligase with reaction progression monitored using LCMS except for protein used in X-ray crystallography studies. Nucleotide exchange was necessary to remove GDP and replaced with it non-hydrolyzable GTP analogue GMPPNP (Jena Bioscience). A Superdex-75 26/600 size-exclusion column (Cytiva) was used for the final purification step in 25 mM HEPES pH 7.5, 150 mM NaCl, 2 mM TCEP, 2 mM MgCl_2_. Monodispersed peak fractions were pooled and concentrated to 2 mg mL^−1^ for BioNMR, biophysical and biochemical characterization activities or 10 mg mL^−1^ for crystallization experiments.

### Surface Plasmon Resonance (SPR)

SPR binding experiments were performed on a Biacore™ S200 instrument (Cytiva) using the Biotin CAPture Kit and Series S Sensor Chip CAP (Cytiva) according to manufacturer protocol. Biotin-labeled RAS proteins were captured using the Biotin CAPture reagent, which consists of streptavidin conjugated to single-stranded DNA that hybridizes to complementary DNA immobilized on the sensor chip, enabling surface regeneration between assay cycles.

Biotin-RAS proteins (D-KRAS4B^G12D^, KRAS4B^WT^, KRAS4B^G12D^, KRAS4B^G12V^, NRAS^WT^, or HRAS^WT^) were prepared at 0.5 µg/mL in assay buffer containing 10 mM HEPES (pH 7.4), 150 mM NaCl, 5 mM MgCl₂, and 0.005% Tween-20, supplemented with 10 µM nucleotide (GDP, GTP, or GMPPCP) corresponding to the RAS nucleotide state. Peptide stocks (10 mM in DMSO) were acoustically dispensed into 384-well plates using an Echo 655T liquid handler and backfilled with DMSO to 2 µL, followed by addition of 98 µL assay buffer using a Bravo liquid handler to yield a final volume of 100 µL and 2% DMSO. Peptides were tested as a five-point, three-fold serial dilution with a top concentration of 1 µM. The SPR running buffer consisted of assay buffer supplemented with 2% DMSO.

All measurements were performed at 25 °C. Sensor chips were primed with running buffer and conditioned with three 60 s injections of regeneration solution (6 M guanidine-HCl, 0.25 M NaOH). Biotin CAPture reagent was injected for 300 s at 2 µL/min, followed by immobilization of biotinylated Ras protein to 300–700 response units (RU) at 10 µL/min flow rate. After a 60 s stabilization period, peptide binding was assessed using single-cycle kinetics with a 120 s association phase (at 20 µL/min flow rate) and a 900 s dissociation phase. Sensor chip surface was regenerated using a 120 s injection of regeneration solution at 10 µL/min before the next cycle. A solvent correction step (using an eight-point [0–3%] DMSO standard) was included to correct for refractive index effects from possible variation in DMSO concentrations. SPR data reported in Extended Data Figure 2 were collected as described above, but without solvent correction, as analyte stocks were prepared in running buffer not containing DMSO. Data were collected using Biacore™ Control software and analyzed with Biacore™ Evaluation software using a 1:1 binding model to calculate rates of association (*k*_on_), dissociation (*k*_off_), and equilibrium dissociation constant (KD).

### Guanine nucleotide exchange (GNE) assay

SOS-mediated nucleotide exchange was assayed as previously reported, with slight procedural modifications. Biotin-labeled RAS protein (2 μM) was treated with EDTA buffer (20 mM HEPES, pH 7.4, 50 mM NaCl, 10 mM EDTA, 0.01% w/v Tween 20) for 1 h at room temperature. Protein was then loaded with fluorescently labeled nucleotide by treating the above solution with a 10-fold molar excess of BODIPY-GDP in SOS assay buffer (20 mM HEPES, pH 7.4, 50 mM NaCl, 10 mM MgCl_2_, 0.01% w/v Tween 20) for 6 h at room temperature. A 1.5X mixture comprising 1.5 nM RAS protein, 15 nM BODIPY-GDP, and 0.25 nM Tb-streptavidin was prepared. Analyte solutions in DMSO were echo-dispensed to a dry plate (Corning3820) before 6 μL of RAS protein solution was added. The plate was sealed and incubated at room temperature for 60 min prior to the addition of 3 μL of 120 nM SOS1 (AA564-1049) and 9 mM GTP premixture. Plates were read after incubation at room temperature for 60 min using an EnVision plate reader with excitation at 340 nm and emission at 495 and 520 nm. Dose-response curves a were analyzed and half-maximal effect concentrations computed using a 4-parameter logistic equation in GraphPad Prism software (GraphPad, San Diego, CA).

### CRAF RBD–KRAS^G12D^(ON) FRET assay

Disruption of the interaction between KRAS^G12D^(ON) and CRAF RBD was assessed biochemically using biotinylated KRAS4B^G12D^ (residues 1-169) loaded with GMPPCP and CRAF (residues 52-131) bearing an N-terminal GST tag. Specifically, 10 nL of test peptide is dispensed into an assay plate (Corning, catalog #3820) using an Echo Acoustic Liquid Handler to make a 10-point, ∼3-fold dilution. To the peptide assay plate, 5 μL of a 2X solution of biotinylated KRAS^G12D^ protein (GMPPCP-loaded; prepared in 20 mM HEPES pH 7.5, 150 mM NaCl, 10 mM MgCl_2_, 0.01% Tween-20) was added and the mixture was incubated at ambient temperature for 30 minutes. Next, a 2X solution of GST-tagged CRAF RBD, anti-GST-d2 acceptor (Cisbio catalog #61GSTDLA) and Streptavidin-Tb cryptate donor (Cisbio catalog #610SATLA) prepared in the same assay buffer was added to a final volume of 10 μL. The final concentrations in the assay were: 10 nM KRAS^G12D^(GMPPCP); 50 nM GST-CRAF RBD; 2 nM anti-GST-d2; and 25 nM SA-Tb cryptate. The assay plate was incubated at ambient temperature with gentle shaking for 60 minutes to allow the reaction to achieve equilibrium. Time-resolved fluorescence resonance energy transfer signal was measured on an Envision (PerkinElmer) plate reader with the following settings: dichroic mirror = LANCE/DELFIA DUAL/Bias; Emission1 = 615 nm; Emission 2 = 665 nM; delay time = 60 ms. The TR-FRET signal from each well was determined as the ratio of the emission at 665 nm to that at 615 nm. Percent effect at each well was determined after normalization to control wells containing DMSO (no effect) or a saturating concentration of an antagonist control peptide (maximum effect). The percent effect as a function of compound concentration was fit to a four-parameter logistic equation.

### CETSA

Macrocyclic peptide cellular target engagement to KRAS^G12D^(GDP) was characterized using HiBIT CETSA as previously described.^24^ MiaPaCa-2 HiBIT-tagged KRAS^G12C^ line was purchased from Promega. MiaPaCa-2 HiBIT (homozygous KRAS^G12C^) cells were seeded in T75 flasks (0.5 million cells/15 ml in Dulbecco’s Modified Eagle’s Medium (DMEM) supplemented with 10% Fetal Bovine Serum (FBS), 2.5% horse serum). The following day, cells were treated with peptides for 2 hr and then trypsinized (0.25%) and pelleted after spinning at 300 rpm for 5 min. For experiments with digitonin treatment, cells were pre-incubated with 10 µg mL^−1^ digitonin for 10 mins before compound dosing. After a PBS wash, the cell pellets were resuspended with PBS (30 µL/well/dose concentration) containing protease inhibitor (Nacalai Tesque) and transferred into 96 well PCR plate (Biorad) and then heated at temperature gradient between 45 to 75 °C for 3 min followed by cooling to 25 °C for 3 min using eppendorf Mastercycler X50a PCR machine. After this step, the cells were frozen immediately in liquid nitrogen and stored at −80 °C. The cells were then thawed and 30µL of 2X stock lysis buffer (100 mM HEPES (pH 7.5), 10 mM beta-glycerophosphate, 0.2 mM sodium orthovanadate, 20 mM MgCl_2_, 4 mM TCEP, 0.05% of w/v Rapigest (Waters) and protease inhibitor (Nacalai Tesque)) was added to each PCR well. TCEP and Rapigest were prepared freshly. After the addition of lysis buffer, the cells were frozen and thawed three times in liquid nitrogen and were transferred to 1.5 mL Eppendorf tubes and were spun at 21,000 *g* and 4 °C for 20 min. Next, 50 µL of supernatant was transferred to a 96 well white bottomed plate (Greiner - 655098) and 50 µL of Nano-Glo HiBIT lytic reagents (Nano-Glo® HiBiT Lytic Detection System-N3030) prepared in 1X lysis buffer was added. Luminescence was measured using Envision Xcite Multilabel Reader (PerkinElmer 1040900). Dose response curves using a 4-parameter logistic equation in GraphPad Prism software (GraphPad, San Diego, CA).

### KRAS^G12D^ cell lysate membrane-patch pull-down

Membrane patches were prepared from AsPC-1 cells using the Minute™ Plasma Membrane Protein Isolation and Cell Fractionation Kit (Invent Biotechnologies SM-005) according to the manufacturer’s protocol. Pellet was resuspended in KRAS IP buffer (PBS pH 7.4, 1 mM MgCl_2_, 1% DMSO, 1X protease inhibitor, 1 mM PMSF) and sonicated briefly. Protein concentration was determined using the BCA protein assay kit (Pierce). Streptavidin beads (10 μL, CST #5947) was mixed with 100 μM biotinylated peptides in 100 μL KRAS peptide buffer (PBS pH 7.4, 1 mM MgCl_2_, 5% DMSO) and incubated on ice for 15 min. Beads were washed two times in ice-cold KRAS peptide buffer. Membrane patches (50 μg) were added together with DMSO or untagged peptides at the indicated concentrations and incubated on ice for 30 min. Beads were washed four times in ice-cold KRAS peptide buffer. After the final wash, bound proteins were eluted with 1X Bolt™ LDS sample buffer supplemented with NuPAGE^®^ sample reducing agent and resolved using Western blot. Primary antibodies used were: RAS (G12D mutant specific, CST #14429) and actin (Santa Cruz sc-1616).

### BRAF–HA-KRAS^G12D^ co-immunoprecipitation

HEK293 cells expressing HA-tagged KRAS^G12D^ were lysed in ice-cold cell lysis-buffer (Cell Signaling Technology) supplemented with 1 mM PMSF and cOmplete™ EDTA-free protease inhibitor cocktail (Roche) for 30 min with intermittent vortexing. Lysates were centrifuged at 18,000 *g* at 4 °C for 15 min and supernatants were then snap-frozen in liquid nitrogen. Protein concentration was determined using the BCA protein assay kit (Pierce). Protein (200 μg) was mixed with 10 μL anti-HA magnetic beads (Cell Signaling Technology, #11846) in the presence or absence of peptides, and the resulting suspensions were incubated overnight at 4 °C. Beads were then washed three times with ice-cold cell-lysis buffer. After the final wash, bound proteins were eluted with 1X Bolt™ LDS sample buffer supplemented with NuPAGE^®^ sample reducing agent and resolved using Western blot as described above. Primary antibodies used were HA-tag (Cell Signaling Technology #2367) and BRAF (Santa Cruz Biotechnology sc-5284).

### ERK phosphorylation AlphaLISA assay

Erk phosphorylation assays was performed as previously reported.^23^ Cellular KRAS inhibitory activity was evaluated by phosphorylation levels of ERK1/2 in AsPC-1 (ATCC® CRL-1682TM, homozygous KRAS^G12D^), Panc 08.13 (ATCC® CRL-2551TM, homozygous KRAS^G12D^), SK-CO-1 (ATCC® HTB-39TM, heterozygous KRAS^G12V^) and A-375 (ATCC® CRL-1619TM, homozygous BRAF^V600E^) cells. AsPC-1 and Panc 08.13 cells were cultured in T175 flask in growth medium (RPMI 1640 Medium, GlutaMAX™ Supplement, HEPES [Gibco 72400-047] supplemented with 10% fetal bovine serum [Hyclone SH30071.03] and 1X Penicillin/Streptomycin [Gibco 15140-122]). Panc 08.13 was further supplemented with 20 μg mL^−1^ of human recombinant insulin (Gibco 12585-014). SK-CO-1 cells were cultured in MEM Medium GlutaMAX™ Supplement (Gibco 42360-032) supplemented with 10% fetal bovine serum (Hyclone SH30071.03) and 1X Penicillin/Streptomycin (Gibco 15140-122). A-375 cells were cultured in (DMEM Medium, GlutaMAX™ Supplement [Gibco 10566-016] supplemented with 10% fetal bovine serum [Hyclone SH30071.03] and 1X Penicillin/Streptomycin [Gibco 15140-122]).

Cells were harvested after 5 min of 0.25% Trypsin-EDTA (Gibco 25200-056) digestion and were seeded in 384-well tissue culture treated plate (Greiner 781091) with respective growth media at a density of 15,000 cells/20μL/well, and incubated at 37°C, 5% CO2 overnight. A-375 cells were seeded at a density of 10,000 cells/20μL/well. Prior to dosing, seeding medium was removed using the BlueCatBio Bluewasher system and replaced with 20 μL of assay medium without fetal bovine serum (RPMI 1640 Medium, no phenol red (Gibco 11835-030) supplemented with 25 mM HEPES (Gibco 15630-080) and 1X Penicillin/Streptomycin (Gibco 15140-122). Compound dose-response titrations were prepared, and appropriate amounts of compounds were dispensed into the 384-well cell culture assay plate using the Echo 550 liquid handler. 25 μL assay medium was added to achieve a final assay volume of 45 μL. Assay plate was incubated at 37 °C, 5% CO_2_ for 2 hours. After treatment, medium was removed from the plate using the BlueCatBio Bluewasher system, and cells were washed once with 25μL 1X DPBS (Gibco 14190-144). Cells were lysed in 20 μL 1X lysis buffer from Alpha SureFire® Ultra™ Multiplex pERK and total ERK assay kit (PerkinElmer MPSU-PTERK) containing EDTA-free Protease inhibitor cocktail (Roche 11836170001) at ambient temperature with constant shaking at 300 rpm for 10-15 min. Cell lysates were mixed for 10 cycles using the Agilent Bravo 384ST liquid handler system before 10 μL was transferred to OptiPlate-384 plate (PerkinElmer 6007680). Phosphorylated ERK and total ERK levels were detected by Alpha SureFire Ultra Multiplex pERK kit (PerkinElmer MPSU-PTERK) using 5 μL acceptor bead mix and 5 μL donor bead mix, both prepared following the manufacturer’s protocol. Plates were sealed using aluminum sealing tape (Costar 07-200-683) during incubation at ambient temperature with constant shaking at 300 rpm for 1 h (both acceptor and donor). Assay plates were read on a Envision Xcite Multilabel Reader (PerkinElmer 1040900) at ambient temperature, with emission at 535 nm (Total ERK) and emission at 615 nm (Phospho ERK). Ratio of pERK vs. total ERK in each well was used as the final readout. Dose response curves and EC_50_ were analyzed using a 4-parameter logistic equation in GraphPad Prism software (GraphPad, San Diego, CA).

For experiments with digitonin treatment, cells were pre-incubated with 10µg mL^−1^ digitonin for 10 mins before compound dosing. pERK measurements were subsequently performed as described above.

### Cell homogenate stability

Ice-cold, suspended HeLa cells (1 million cells mL^−1^) were homogenized using a probe sonicator, and the resulting homogenate was stored frozen at −20 °C until use. Reactions were prepared as 1 μM solutions of peptide analyte (**DMP-1**) in 50 μL of cell homogenate, and were aged at 37 °C. At 0, 10, 30, 60, and 120 min, reactions were quenched with the addition of 80% v/v acetonitrile in methanol (150 μL) containing a known internal standard. The quenched samples were centrifuged at 4000 rpm for 5 min at 10 °C, and supernatants were analyzed by LCMS. As a negative control, deactivated homogenate (prepared by incubating homogenate at 100 °C for 30 min) was used. The first-order rate constant *k*_e_ describing the reduction in parent peptide concentration over time was computed by regression analysis, and the resulting half-life was calculated as follows: *t*_½_ = ln2 • *k*_e_^−1^.

### NMR spectroscopy

NMR experiments were carried out on a Bruker Avance III HD 800 MHz spectrometer equipped with 5-mm TCI cryoprobe. The two NMR samples are 0.225mM ^13^C/^15^N labeled KRAS^G12D^(GDP), with or without 0.25 mM **DMP-1** peptide, in a 50 mM HEPES, pH 7.4, 50 mM NaCl, 2.0 mM MgCl_2_, 50 µM 3-(trimethylsilyl) propionic-2,2,3,3-*d*4 acid (TSP), and 10% (v/v) D_2_O buffer. All 3D experiments were collected at 300K with nonuniform sampling (NUS) at a sampling rate of 25%. All NMR data were processed on a Linux station using NMRPipe and the spectra were analyzed using NMRFAM-Sparky^36^. The backbone resonance assignment of KRAS^G12D^(GDP) was achieved previous^37^ and was calibrated using triple resonance HNCA and HN(CO)CA. The chemical shift perturbations induced by the peptide binding were calculated using the following formula: Δδ(**DMP-1**) = (((Δδ_1H_)^2^ + (0.14 × Δδ_15N_)^2^) / 2) ^1/2^.

### X-ray crystallography

Purified KRAS^G12D^(GMPPNP) was mixed with a 3-fold molar excess of **DMP-1** and incubated on ice for two hours. Sitting-drop vapor diffusion crystallization screening was conducted by mixing 0.5 μL of protein solution with 0.5 μL of reservoir solution and equilibrating the drops against 100 uL of reservoir solution at 18°C. Crystals suitable for diffraction experiments were obtained from Hampton Research PEG/Ion HT screen condition D2, which consisted of 20% w/v PEG 3350 and 0.2 M ammonium tartrate. A single crystal was harvested from the sitting drop plate and briefly cryoprotected in mother liquor supplemented with 10% glycerol prior to plunge freezing in liquid nitrogen. Diffraction data were collected at Advanced Photon Source IMCA-CAT beamline 17-ID and processed using AutoPROC.^38^ The structure was solved by molecular replacement using an internal structure of monomeric KRAS^G12D^ as a search probe. Ligand restraints were generated using CCP4 AceDRG.^39^ Manual ligand fitting with subsequent rounds of model building and iterative refinement were conducted in Coot^40^ and BUSTER.^41^ The final coordinates and structure factors were deposited in the Protein Data Bank under accession code 12XR. A summary of the diffraction and refinement statistics can be found in Supplementary Table 1.

### Protein FEP simulation

The X-ray co-crystal structure of **DMP-1** and GMPPNP bound to KRAS4B reported here was prepared using the Protein Preparation Wizard in Schrödinger Suite (Schrödinger, LLC; release 2025-2), including assignment of bond orders, addition of hydrogens, optimization of hydrogen-bonding networks, and restrained minimization using the OPLS4 force field. Protonation states of ionizable residues at pH 7.4 were assigned using Epik. The peptide ligand was retained in the binding site as observed in the experimental structure, and was treated as the ligand in subsequent calculations. Relative binding free-energy calculations for D153E mutation were performed using the Protein FEP workflow in FEP+ (Schrödinger Suite 2025-2). The FEP graph contained two nodes (wild-type and D153E mutant) and one edge. For each state (protein-peptide complex, apo protein/solvent, and fragment), Desmond systems were generated. Systems were solvated with explicit SPC water in a periodic μVT ensemble-compatible box, using the default FEP+ box shape and size parameters with 5.0 Å buffer between solute and box edges. Counterions were added to neutralize the system; no additional salts were included in the simulation. The WT→D153E alchemical transformation was decomposed into 12 lambda windows for the neutral mutation. Each window was equilibrated for 0.5 ns, followed by 10 ns of production MD for the complex, solvent, and fragment legs (10/10/10 ns per state). Free energies for each leg were obtained using Bennett’s acceptance ratio as implemented in FEP+, and combined to yield the change in binding free energy predicted for D153E relative to WT (ΔΔG_bind_ = 2.9 ± 0.46 kcal mol^−1^). To assess convergence, an additional Protein FEP calculation was performed with extended production time of 15 ns per lambda window. The resulting ΔΔG_bind_ = 2.9 ± 0.44 kcal mol^−1^ was statistically indistinguishable from the 10 ns-per-window result, supporting the convergence and robustness of the prior result.

