## Supplementary Information for "Mirror-image mRNA display uncovers isoform-selective D-peptide macrocycles targeting a cryptic KRAS pocket"

### X-Ray Crystallography

**Supplementary Table 1: Data collection and refinement statistics**

| **PDB ID: 12XR** | **KRAS$^{G12D}$(GMPPNP) : DMP-1 complex** |
|---|---|
| **Wavelength (Å)** | 1.000 |
| **Resolution range (Å)** | 62.48 - 1.48 (1.50 - 1.48) |
| **Space group** | P 21 21 2 |
| **Unit cell – (a, b, c, α, β, γ)** | 80.36 41.48 62.48 90 90 90 |
| **Total reflections** | 213563 (4502) |
| **Unique reflections** | 33766 (1108) |
| **Multiplicity** | 6.3 (4.1) |
| **Completeness (%)** | 94.5 (63.4) |
| **Mean I/sigma(I)** | 8.7 (0.5) |
| **Wilson B-factor (Å$^2$)** | 21.1 |
| **R-merge** | 0.100 (2.343) |
| **CC1/2** | 0.998 (0.401) |
| **Reflections used in refinement** | 33612 (673) |
| **Reflections used for R-free** | 1659 (33) |
| **R-work** | 0.1908 (0.3789) |
| **R-free** | 0.2136 (0.3651) |
| **Number of non-hydrogen atoms** | 1724 |
| **RMS bonds (Å)** | 0.010 |

| | |
|---|---|
| **RMS angles (°)** | 0.99 |
| **Ramachandran favored (%)** | 97.62 |
| **Ramachandran allowed (%)** | 2.38 |
| **Ramachandran outliers (%)** | 0.00 |
| **Average B-factor (Å$^2$)** | 23.83 |

Statistics for the highest-resolution shell are shown in parentheses.

### Synthesis of macrocyclic peptides

#### General materials and methods

Macrocyclic peptides were synthesized using standard solid-phase synthesis techniques, employing Fmoc/*t*-Bu chemistry as described in Chan, W. C.; White, P. D. "Fmoc Solid-Phase Synthesis: a Practical Approach", Oxford University Press, Oxford, 2000; Steward, J.; Young, J. "Solid Phase Peptide Synthesis", Pierce Chemical Company, Rockford, 1984.; N. L. Benoiton, "Chemistry of Peptide Synthesis", CRC Press, New York, 2006; and Lloyd-Williams, P.; Albericio, F. "Chemical Approaches to the Synthesis of Peptides and Proteins", CRC Press, New York, 1997. Fmoc (9*H*-fluoren-9-ylmethoxycarbonyl) protected monomers were obtained from Sigma-Aldrich, Novabiochem, Chem-Impex, and Combi-Blocks.

#### Synthesis of SII-P ligand mirror-image pair MP-9312 and ent-MP-9312

A macrocyclic peptide ligand of the KRAS switch-II pocket (**MP-9312**), closely related in structure to those previously described,[1] was prepared alongside its mirror-image counterpart (**ent-MP-9312**). Affinity of these ligands to L- and D-KRAS$^{G12D}$ was assessed using TR-FRET and SPR, confirming the competency of D-KRAS$^{G12D}$ to bind mirror-image analogs of established SII-P ligands.

**Supplementary Table 2: Sequence and affinity data for SII-P peptides (enantiomeric pair)**

| Compound | Sequence | L-KRAS$^{G12D}$ SII-P TR-FRET $EC_{50}$ [nM] | D-KRAS$^{G12D}$ SPR $K_D$ [nM] |
|---|---|---|---|
| **MP-9312** | Ac-G-cyclo(DC(methylene)-P-L-Y-I-S-Y-D-P-V-C)-$NH_2$ | 550 | ND |
| ***ent*-MP-9312** | Ac-G-cyclo(C(methylene)-DP-DL-DY-DI-DS-DY-DD-DP-DV-DC)-$NH_2$ | >48,080 | 1.2 |

[1] Lim, S. *et al.* Discovery of cell active macrocyclic peptides with on-target inhibition of KRAS signaling. *Chem. Sci.* **12**, 15975-15987 (2021).

Synthesis of **MP-9312** and ***ent*-MP-9312** was achieved as follows:

***Step 1:*** **Solid phase peptide synthesis.** Linear peptide was synthesized on 50 µmol (**MP-9312**) or 100 µmol (***ent*-MP-9312**) scale using Rink Amide resin LL MBHA (NovaBioChem, 0.33 mmol/g loading) and a standard Fmoc/*t*-Bu.protection schemes on CEM Liberty automated microwave peptide synthesizer. Specifically, *tert*-butyl side-chain protection was employed for Asp, Tyr, Ser or their D-counterparts. Trityl side chain protection was employed for Cys and d-Cys. Single couplings were performed using 5 equiv. Fmoc-AA, 10 equiv. DIC, and 5 equiv. Oxyma Pure at 90 °C for 2 min for Gly, Pro (after Leu), Leu, Tyr, Ile, Ser, Asp residues or their D-counterparts, while double couplings were performed for more difficult couplings such as for D-Cys, Pro (after Val) and Cys residues. One double coupling extended (90 °C, 4 min) was performed for the Val residue. Fmoc removal was achieved before each coupling by treatment with 20% v/v piperidine in DMF at 90 °C for 2 min. Acetylation of the N terminus of the fully assembled linear sequence was performed using acetic anhydride (10% v/v in DMF, 75 °C for 10 min). The resin-supported peptide was then thoroughly washed with DMF and DCM and allowed to dry under reduced pressure.

***Step 2:*** **Cleavage from the resin and isolation of peptide intermediate.** Linear peptide was cleaved from the resin by treatment with 5 mL of a cleavage cocktail comprising 94% TFA, 3% triisopropylsilane, and 3% water at 40 °C for 30 min using a Razor peptide cleavage system from CEM corporation. After filtration from the resin, the cleavage solution was concentrated to a total volume of approximately 2 mL before cold methyl tert-butyl ether (25 mL) was added to precipitate the crude peptide. The suspension was cooled down in dry ice for 60 min. After centrifugation for 15 min, the supernatant was removed. Methyl tert-butyl ether (25 mL) was added to the white pellet. The suspension was cooled in dry ice for 60 min. After centrifugation for 15 min, the supernatant was removed. The white pellet was allowed to air dry.

The residue was then semi-purified by reverse-phase flash column chromatography (C18 stationary phase, eluting with 20% MeCN/water containing 0.1% TFA initially, grading to 60% MeCN/water containing 0.1% TFA). Product fractions were collected and freeze-dried to provide the linear peptide product.

***Step 3:*** **Cyclization and isolation.** The linear peptide was dissolved in 50% v/v acetonitrile/water (25 mL), and the pH of the resulting solution was adjusted to pH ~ 9-10 by addition of N,N-diisopropylethylamine. DL-Dithiothreitol (DTT, 2 equiv.) and diiodomethane (10 equiv.) were then added at room temperature, and acetonitrile (~ 2 mL) was added as needed to attain a homogeneous reaction mixture. This solution was aged at room temperature overnight, whereupon it was acidified with the introduction of TFA (100 μL), concentrated under reduced pressure, and freeze-dried. The residue was purified by reverse-phase HPLC (C18 stationary phase, eluting with a gradient of MeCN in water containing 0.1% TFA). Product fractions were collected and freeze-dried to provide the desired cyclic peptide as a white amorphous solid.

**Supplementary Table 3: Characterization of synthesized peptides**

| Peptide | Purity (%) | Yield (%) | Formula | MW | Exact mass $[M+2H]^{++}$ | Observed mass $[M+2H]^{++}$ |
|---|---|---|---|---|---|---|
| **MP-9312** | 100 | 12 | $C_{63}H_{91}N_{13}O_{18}S_2$ | 1382.6 | 691.8 | 692.3 |
| ***ent*-MP-9312** | 100 | 9 | $C_{63}H_{91}N_{13}O_{18}S_2$ | 1382.6 | 691.8 | 692.0 |

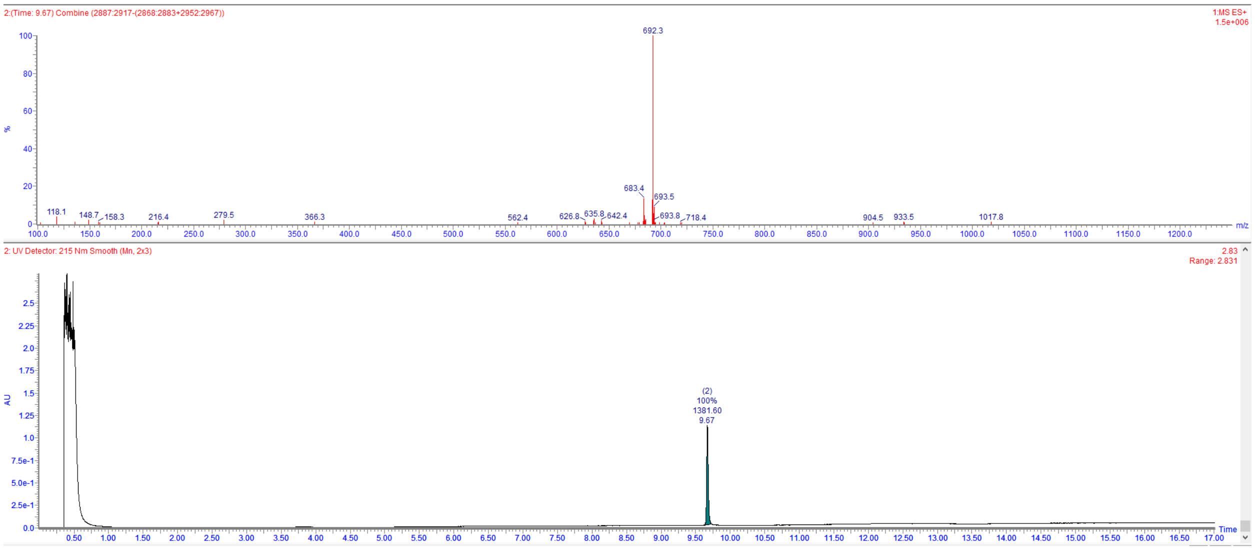

**Supplementary Figure 1: LCMS characterization of MP-9312.**

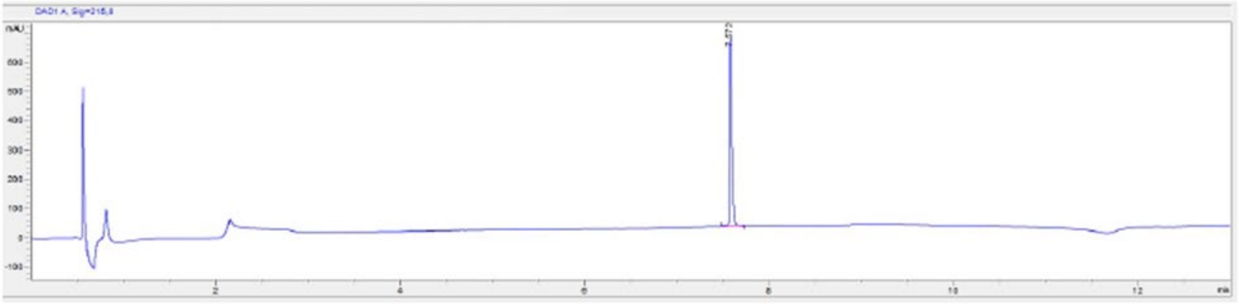

| # | Time | Type | Area | Height | Width | Area% | Symmetry |
|---|---|---|---|---|---|---|---|
| 1 | 7.572 | BB | 1074.5 | 650 | 0.0238 | 100.000 | 0.494 |

Ret. Time: 7.57

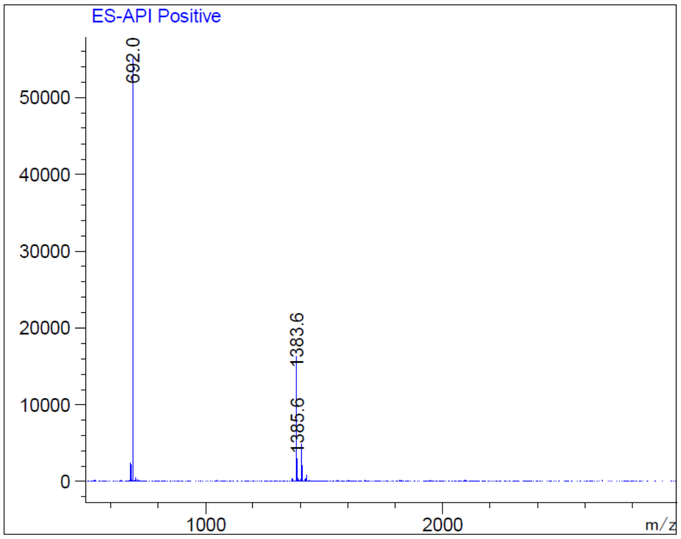

**Supplementary Figure 2: LCMS characterization of ent-MP-9312.**

**Synthesis of ᴅMP-1 and ᴅMP-NBC**

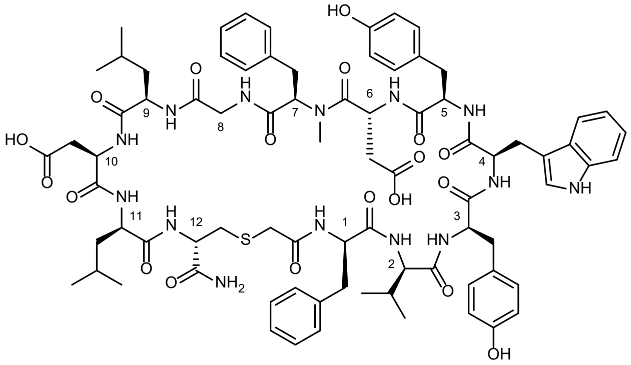

Synthesis of **ᴅMP-1** and **ᴅMP-NBC** was achieved as follows:

***Step 1:*** **Solid phase peptide synthesis.** Linear peptide was synthesized on 100 µmol scale (2 × 50 µmol scale syntheses performed in parallel) using Fmoc-Sieber-PS resin (0.7 mmol/g loading) and a standard Fmoc/*t*-Bu protection scheme on a SyroWave automated synthesizer. Specifically, the side chain protecting groups employed were: *tert*-butyl for ᴅ-Asp and ᴅ-Tyr; trityl for ᴅ-Cys; and Boc for ᴅ-Trp. Single couplings were employed for all residues (5 equiv. Fmoc-AA, 5 equiv. DIPCDI, and 5 equiv. Oxyma Pure in NMP), conducted at 50 °C for 15 min each. Removal of Fmoc protecting groups was achieved through treatment with 20% v/v pyrrolidine in NMP at 50 °C for 15 min. The *N*-terminus of the fully assembled, linear, resin-supported peptide was capped by treatment with chloroacetic anhydride (5 equiv. as a 0.2 M NMP solution) at room temperature for 1 h. The resin-supported peptide was then thoroughly washed with NMP and MeOH, and then dried under reduced pressure.

***Step 2:*** **Cleavage from the resin and cyclization of peptide intermediate.** Linear peptide was cleaved from the resin by treatment with 10 mL of a cleavage cocktail comprising 80% TFA, 5% *m*-cresol, 5% thioanisole, 5% water, 2.5% 3,6-dioxa-1,8-octanedithiol (DODT), and 2.5% triisopropylsilane at room temperature for 90 min. The cleavage solution was then concentrated to a total volume of approximately 1 mL before cold diethyl ether was added to precipitate the peptide product. After centrifugation, the ether was decanted, and the pellet dissolved in approximately 10 mL of 50% acetonitrile in water; this mixture was filtered, and the filtrate was treated with triethylamine (70 µL, 10 equiv.) to perform the intramolecular cyclization. The resulting mixture was aged at room temperature for 1 h. The reaction mixture was then acidified with the addition of AcOH (29 µL, 10 equiv) before it was freeze-dried. The residue thus obtained was purified by reverse-phase HPLC (L-column2 ODS stationary phase, eluting with acetonitrile/water containing 0.1% TFA modifier) to provide pure cyclic peptide as a white solid after lyophilization.

**Supplementary Table 4: Characterization of synthesized peptides**

| Peptide | Purity (%) | Yield (%) | Formula | MW | Exact mass $[M+2H]^{++}$ | Observed mass $[M+2H]^{++}$ |
|---|---|---|---|---|---|---|
| **DMP-1** | 97% | 13% | $C_{80}H_{100}N_{14}O_{19}S$ | 1593.8 | 797.4 | 797.7 |
| **DMP-NBC** | 98% | 17% | $C_{80}H_{100}N_{14}O_{19}S$ | 1593.8 | 797.4 | 797.8 |

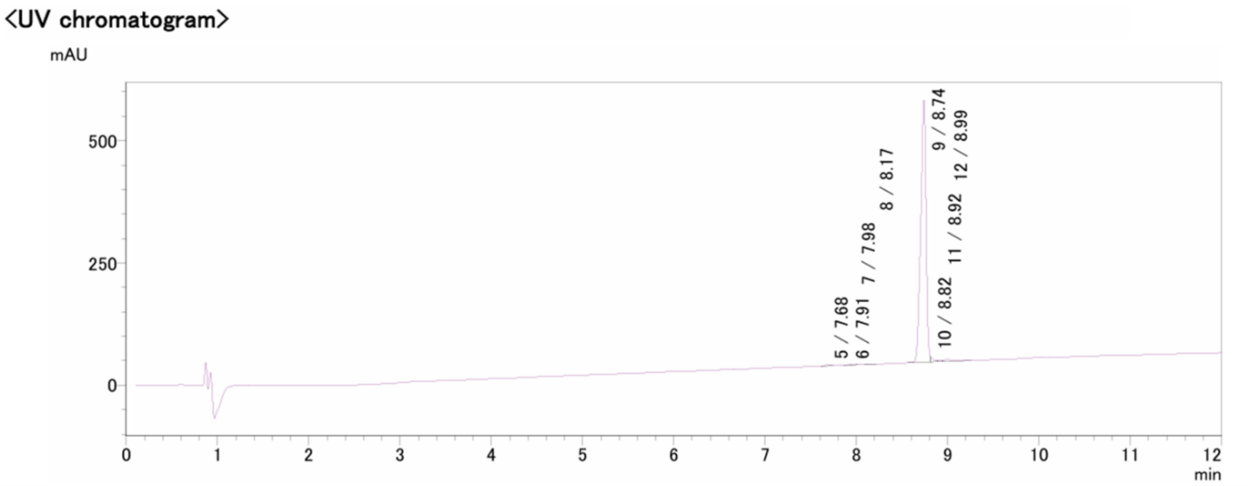

UV peak table

PDA Ch1 220nm

| Peak# | Time | Area | Area% | Height |
|---|---|---|---|---|
| 5 | 7.68 | 8014 | 0.36 | 2269 |
| 6 | 7.91 | 1557 | 0.07 | 544 |
| 7 | 7.98 | 1481 | 0.07 | 522 |
| 8 | 8.17 | 2022 | 0.09 | 559 |
| 9 | 8.74 | 2175088 | 96.94 | 535986 |
| 10 | 8.82 | 23628 | 1.05 | 9950 |
| 11 | 8.92 | 6996 | 0.31 | 2404 |
| 12 | 8.99 | 24882 | 1.11 | 4311 |

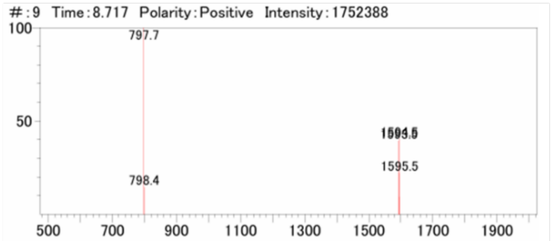

**Supplementary Figure 3: LCMS characterization of DMP-1**

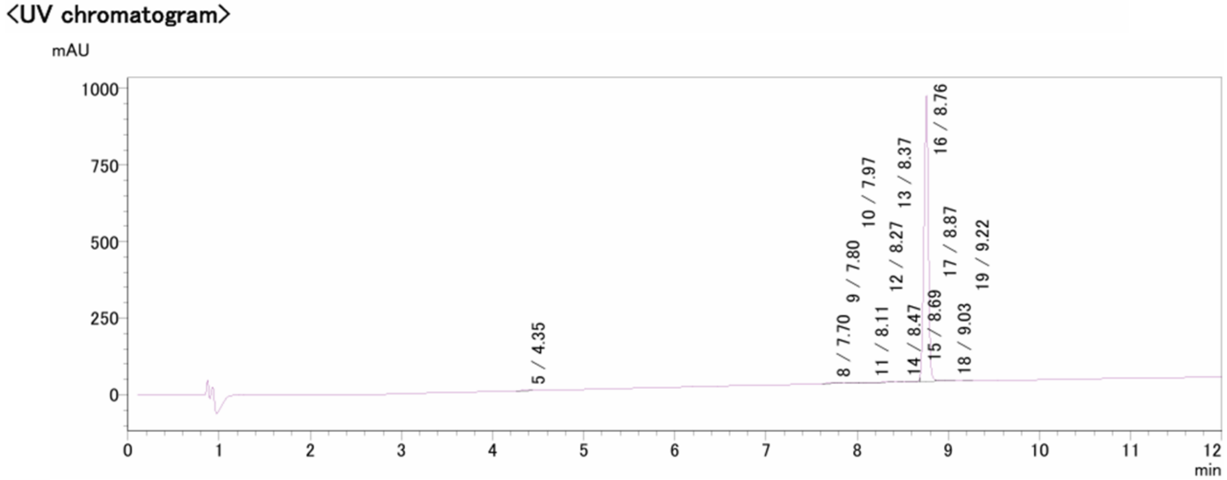

**UV peak table**

PDA Ch1 220nm

| Peak# | Time | Area | Area% | Height |
|---|---|---|---|---|
| 5 | 4.35 | 7428 | 0.24 | 1344 |
| 8 | 7.70 | 10627 | 0.34 | 2643 |
| 9 | 7.80 | 3788 | 0.12 | 1620 |
| 10 | 7.97 | 7000 | 0.22 | 2021 |
| 11 | 8.11 | 3958 | 0.13 | 616 |
| 12 | 8.27 | 573 | 0.02 | 237 |
| 13 | 8.37 | 512 | 0.02 | 194 |
| 14 | 8.47 | 708 | 0.02 | 272 |
| 15 | 8.69 | 7676 | 0.24 | 6660 |
| 16 | 8.76 | 3105749 | 98.31 | 933313 |
| 17 | 8.87 | 7474 | 0.24 | 3506 |
| 18 | 9.03 | 1589 | 0.05 | 656 |
| 19 | 9.22 | 2174 | 0.07 | 751 |

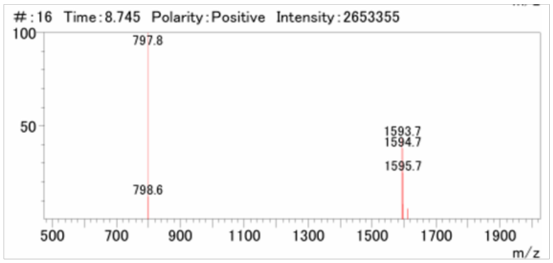

**Supplementary Figure 4: LCMS characterization of** **ᴅMP-NBC**

### Biochemical profiling

#### Synthesis of DMP-1 TR-FRET tracer

KRAS cryptic back-pocket binding was measured by a TR-FRET competition binding assay employing a fluorescein (5-FAM) labeled analog of **DMP-1** with the following structure (**DMP-1-FITC**):

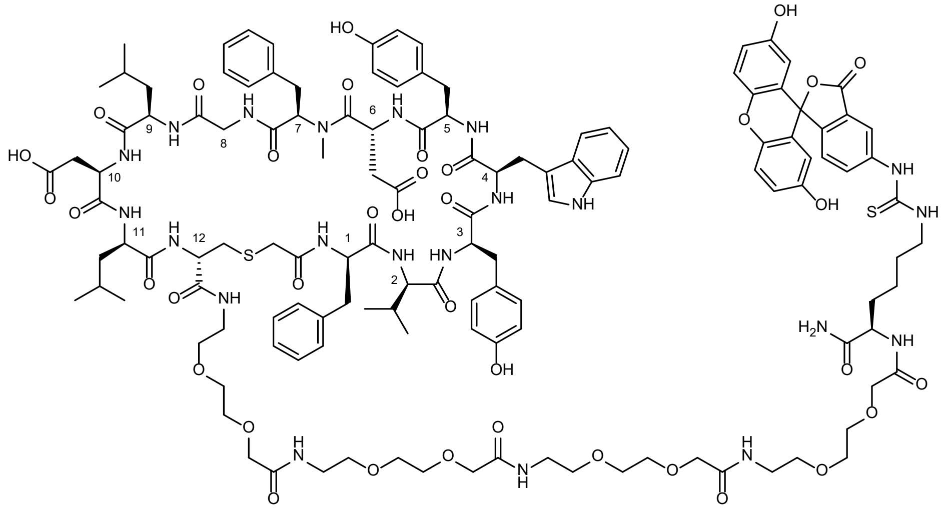

***Step 1:*** **Solid phase peptide synthesis.** Linear peptide was synthesized on 100 µmol scale using Rink amide MBHA resin (EMD corp., 0.5 mmol/g loading) and a standard Fmoc/*t*-Bu protection schemes on a Symphony X automated synthesizer. Specifically, the side chain protecting groups employed were: *tert*-butyl for D-Asp, D-Tyr, and D-Cys; and Boc for D-Lys and D-Trp. Single couplings were employed for all residues (8 equiv. Fmoc-AA as 0.2 M DMF solution, 8 equiv. HATU as 0.4 M DMF solution, and 16 equiv. NMM as 0.8 M DMF solution). Amino acids with β-branching (D-Val2) and amino acids coupled to *N*-methylated residues (D-Asp6) were coupled for 60 min, whereas other couplings were performed for 10 min. Removal of Fmoc protecting groups was achieved through three cycles of treatment with 20% v/v pyrrolidine in DMF, each cycle lasting 5 minutes. The N terminus of the fully assembled, linear, resin-supported peptide was capped by treatment with chloroacetic anhydride (10 equiv. as a 0.2 M DMF solution), lasting 15 minutes. The resin-supported peptide was then thoroughly washed with DMF and DCM and allowed to dry under reduced pressure.

***Step 2:*** **Cleavage from the resin and cyclization of peptide intermediate.** Linear peptide was cleaved from the resin by treatment with 10 mL of a cleavage cocktail comprising 95% TFA, 2.5%

triisopropylsilane, and 2.5% water at room temperature. The cleavage solution was then concentrated to a total volume of approximately 1 mL before diethyl ether was added and the diluted ethereal mixture was cooled on dry ice for 10 minutes in order to precipitate the peptide product. The chilled mixture was centrifuged (5,000 rpm, 30 min), the ether decanted, and the pellet dissolved in approximately 10 mL of 50% acetonitrile in water; pH was adjusted to ~8 by the addition of 0.2 M aqueous ammonium bicarbonate solution. Thioether macrocyclization was confirmed by LCMS analysis before the solution was freeze-dried to provide crude macrocyclic peptide intermediate. This substance was purified by reverse-phase HPLC using a C8 column with acetonitrile/water containing 0.1% TFA modifier to provide pure cyclic peptide intermediate (35 mg, 92% purity, 14% yield). MS: Expected *m/z* 1152.6, observed 1152.6 $[M+2H]^{++}$.

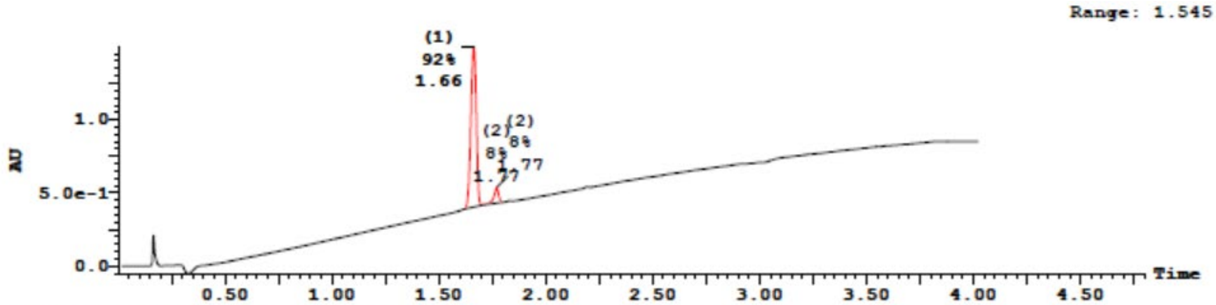

| Peak Number | Compound | Time | AreaAbs | Area %Total | Width | Height | Mass Found |
|---|---|---|---|---|---|---|---|
| 1 | | 1.66 | 3e+004 | 92.04 | 0 | 1e+006 | Not Found |
| 2 | | 1.77 | 3e+003 | 7.96 | 0 | 1e+005 | Not Found |

| Peak ID | Compound | Time | Mass Found |
|---|---|---|---|
| 1 | | 1.66 | Not Found |

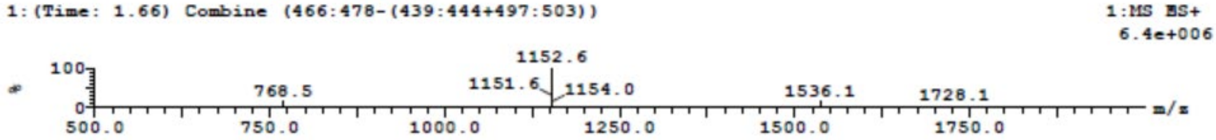

**Supplementary Figure 5. LCMS characterization of unlabeled precursor to DMP-1-FITC**

***Step 3:*** **Fluorophore installation and isolation of FRET tracer.** Peptide intermediate bearing a C-terminal deprotected D-Lys functional handle (25 mg, 11 μmol) was dissolved in DMF (2 mL). Fluorescein 5-isothiocyanate (8.5 mg, 22 μmol, 2 equiv.) and *N*,*N*-diisopropylethylamine (8.9 μL, 54 μmol, 5 equiv.) were added sequentially at room temperature. The reaction stirred for 1 h, whereupon LCMS analysis confirmed complete conversion to the desired product. The mixture was purified by reverse-phase HPLC using a C8 column with acetonitrile/water containing 0.1%

TFA modifier to provide pure **DMP-1-FITC** (9.0 mg, 88% purity, 28% yield). MS: Expected *m/z* 1346.1, observed 1347.2 $[M+2H]^{++}$.

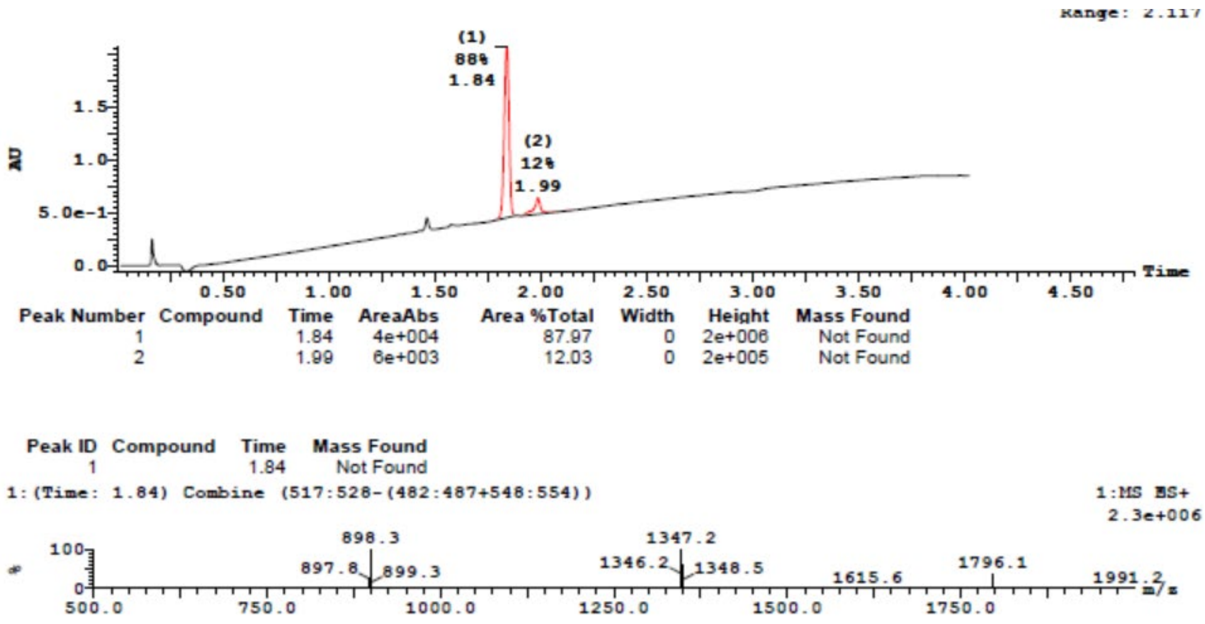

**Supplementary Figure 6. LCMS characterization of DMP-1-FITC**

### Time-Resolved Fluorescence/Förster Resonance Energy Transfer (TR-FRET)

Time-Resolved Fluorescence/Förster Resonance Energy Transfer (TR-FRET) Assay using KRAS G12D or KRAS G12V loaded with GDP or GMP-PCP: Macrocyclic peptide binding affinities to KRAS variants were characterized using a TR-FRET–based competitive binding assay. Compound dose-response titrations were prepared, and appropriate amounts of compounds were dispensed, using Echo 550 liquid handler, into the 384-well black microplate (Corning 3820). A solution of 1.33x KRAS or buffer without KRAS was added to the assay plate and incubated with compounds for 60 minutes. A solution of 4x Terbium-Streptavidin and fluorescently labeled tracer (**DMP-1-FITC**) was prepared and added to assay plate with final assay volume of 10µL. Assay plates were read at ambient temperature on the Envision, with excitation at 340 nm and emission at 495 and 520 nm. Dose response curves and EC50 were analyzed using a 4-parameter logistic equation in GraphPad Prism software (GraphPad, San Diego, CA).

### Sequence alignment of RAS family and related GTPases

Uniprot sequences for KRAS4A (P-01116-1), KRAS4B (P-01116-2), HRAS4A (P01112-1), HRASIDX (P01112-2), and NRAS (P01111) were processed by the T-Coffee Expresso sequence alignment tool.[2,3] Whole-protein alignments are as follows:

```
T-COFFEE, Version_11.00 (Version_11.00)
Cedric Notredame

KRAS4A    MTEYKLVVVGAGGVGKSALTIQLIQNHFVDEYDPTIEDSYRKQVVIDGETCLLDILDTAGQEEYSAMRDQYM
KRAS4B    MTEYKLVVVGAGGVGKSALTIQLIQNHFVDEYDPTIEDSYRKQVVIDGETCLLDILDTAGQEEYSAMRDQYM
HRAS4A    MTEYKLVVVGAGGVGKSALTIQLIQNHFVDEYDPTIEDSYRKQVVIDGETCLLDILDTAGQEEYSAMRDQYM
HRASIDX   MTEYKLVVVGAGGVGKSALTIQLIQNHFVDEYDPTIEDSYRKQVVIDGETCLLDILDTAGQEEYSAMRDQYM
NRAS      MTEYKLVVVGAGGVGKSALTIQLIQNHFVDEYDPTIEDSYRKQVVIDGETCLLDILDTAGQEEYSAMRDQYM

cons      ************************************************************************

KRAS4A    RTGEGFLCVFAINNTKSFEDIHHYREQIKRVKDSEDVPMVLVGNKCDLPSRTVDTKQAQDLARSYGIPFIET
KRAS4B    RTGEGFLCVFAINNTKSFEDIHHYREQIKRVKDSEDVPMVLVGNKCDLPSRTVDTKQAQDLARSYGIPFIET
HRAS4A    RTGEGFLCVFAINNTKSFEDIHQYREQIKRVKDSDDVPMVLVGNKCDLAARTVESRQAQDLARSYGIPYIET
HRASIDX   RTGEGFLCVFAINNTKSFEDIHQYREQIKRVKDSDDVPMVLVGNKCDLAARTVESRQAQDLARSYGIPYIET
NRAS      RTGEGFLCVFAINNSKSFADINLYREQIKRVKDSDDVPMVLVGNKCDLPTRTVDTKQAHELAKSYGIPFIET

cons      **************:*** **: ***********:*************.:***:::**::**:*****:***

KRAS4A    SAKTRQRVEDAFYTLVREIR--QYRLKKISKEEKTPGCVKIKKCIIM-------
KRAS4B    SAKTRQGVDDAFYTLVREIR--KHK------EKMSKDGKKKKKKSKTKCVIM--
HRAS4A    SAKTRQGVEDAFYTLVREIRQ--HK------L-RKLNPPDESGPGCMSCKCVLS
HRASIDX   SAKTRQGSRSGSSSSSGTLWDPPGPM----------------------------
NRAS      SAKTRQGVEDAFYTLVREIRQ--YR------M-KKLNSSDDGTQGCMGLPCVVM

cons      ******   ..  :    :
```

[2] Notredame, C., Higgins D.G., Heringa J. T-Coffee: A novel method for fast and accurate multiple sequence alignment. J. Mol. Biol. 302, 205-217 (2000).

[3] Armougom, F. et al. Expresso: automatic incorporation of structural information in multiple sequence alignments using 3D-Coffee. Nucleic Acids Res. 34, W604-W608 (2006).
